# Expanding the Substrate Scope of *N*- and *O*-Methyltransferases from Plants for Chemoselective Alkylation

**DOI:** 10.1101/2023.07.21.549995

**Authors:** Emely Jockmann, Fabiana Subrizi, Michael K. F. Mohr, Eve M. Carter, Pia M. Hebecker, Désirée Popadić, Helen C. Hailes, Jennifer N. Andexer

## Abstract

Methylation reactions are of significant interest when generating pharmaceutically active molecules and building blocks for other applications. Synthetic methylating reagents are often toxic and unselective due to their high reactivity. *S*-Adenosyl-L-methionine (SAM)-dependent methyltransferases (MTs) present a chemoselective and environmentally friendly alternative. The anthranilate *N*-MT from *Ruta graveolens* (*Rg*ANMT) is involved in acridone alkaloid biosynthesis, methylating anthranilate. Although it is known to methylate substrates only at the *N-*position, the closest relatives with respect to amino acid sequence similarities of over 60% are *O*-MTs catalysing the methylation reaction of caffeate and derivatives containing only hydroxyl groups (CaOMTs). In this study, we investigated the substrate range of *Rg*ANMT and a CaOMT from *Prunus persica* (*Pp*CaOMT) using compounds with both, an amino- and hydroxyl group (aminophenols) as possible methyl group acceptors. For both enzymes, the reaction was highly chemoselective. Furthermore, generating cofactor derivatives in situ enabled the transfer of other alkyl chains onto the aminophenols, leading to an enlarged pool of products. Selected MT reactions were performed at a preparative biocatalytic scale in in vitro and in vivo experiments resulting in yields of up to 62%.

## Introduction

*S*-Adenosyl-L-methionine (SAM) is an ubiquitous cofactor.^[1]^ In nature, only 5’-adenosine triphosphate (ATP) is used more often as an enzyme substrate.^[2]^ The chemical structure of SAM was determined in 1952 by *Cantoni et. al*. and can be divided into two main components: an amino acid part arising from L-methionine, and an adenosyl part derived from ATP. Both moieties are linked by the positively charged sulfonium, activating the cofactor for nucleophilic attack.^[3–5]^ In addition, SAM can also act as a source for radicals and as an ylide.^[1,6]^ Methylation is one important function of SAM – the acceptor molecules vary from small compounds such as dopamine or anthranilate (**1**) to macromolecules such as proteins or DNA.^[7–9]^ Under physiological conditions, SAM is unstable and degrades non-enzymatically.^[10,11]^ In the last few years, different multienzyme systems have been established for SAM supply and regeneration.^[12–15]^ In this work we use a three-enzymes cascade as described by others and us:^[16,17]^ ATP and L-methionine are used for the in situ synthesis of the cofactor SAM, catalysed by an L-methionine adenosyltransferase (MAT, EC 2.5.1.6, first step); the methyl group of SAM is transferred onto a substrate by a methyltransferase (MT, EC 2.1.1.x, second step); *S*-adenosyl-L-homocysteine (SAH), the by-product of the previous reaction is then cleaved into adenine and *S*-ribosyl-L-homocysteine by a methyl thioadenosine/ SAH nucleosidase (MTAN, EC 3.2.2.9, third step). Such enzyme cascades offer several advantages: the in situ synthesis ensures a stereoselective production of the cofactor; moreover, the starting substrates (ATP and L-methionine) can be modified to produce more stable derivates.^[18]^ Also, MAT enzymes accept L-methionine analogues forming cofactor derivatives with altered residues such as ethyl, propargyl, allyl and benzyl groups that are further transferred by MTs.^[19– 22]^ Furthermore, SAH, the by-product of the alkylation reaction is known to be an inhibitor for many MTs.^[23]^ Cleavage of SAH catalysed by the third enzyme in the cascade - the MTAN - is irreversible, shifting the equilibrium towards the product side and preventing inhibition effects on the methylation step.

The chemical and physical properties of molecules are altered in different ways by the installation of the methyl group. Not only in nature, but also in pharmaceutical and biotechnological industry, methylation is of great interest. Many small molecule drugs contain at least one methyl group availing the so-called *“magic methyl effect”*.^[24]^ Traditionally, methylation is carried out with reagents such as methyl iodide. Enzyme cascades present a non-toxic and environmentally friendly approach that can be used at scale following process optimisation.^[25]^

MTs are highly chemo- and stereoselective,^[26,27]^ they can be grouped into *C*-, *N*-, *O*-, *S*- and halide MTs, depending on the heteroatom receiving the methyl group. *O*-MTs are the largest group and involved in important reactions such as neurotransmitter deactivation and lignin biosynthesis.^[7,28,29]^ Catechol *O-*MTs (COMTs) are known for the *O-*methylation of dopamine. In previous studies it was found that the COMT from *Myxococcus xanthus (Mx*SafC) was also able to perform a double methylation of tetrahydroisoqunolines while the related enzyme from *Rattus norvegicus* (*Rn*COMT) transferred only one methyl group regioselectively.^[30]^ Different classification systems have been suggested to distinguish between similar enzymes. Regarding plant *O*-MT sequences, *Joshi et al*. introduced a classification system dividing *O*-MTs into class I and II, according to their amino acid sequence, size and dependence on metal ions.^[31]^ Class I *O*-MTs range between 231 and 248 amino acids and have metal ions such as Mg^2+^ as cofactor; this classification can be extended to enzymes from other organisms, grouping mammalian metal ion dependent COMTs such as *Rn*COMT and the bacterial *Mx*SafC into class I. Class II enzymes are typically comprised of 344 – 384 amino acids and are independent of metal ions; a prominent example are class II *O*-MTs catalysing the methylation reaction of caffeate (**2**) (CaOMTs, EC 2.1.1.68). This enzyme family is involved in the biosynthesis of lignin (Figure 1). Besides **2** and derivatives, CaOMT has also shown activity towards 5-hydroxyferulic acid and analogues and methylates at the hydroxyl group at the 3-or 5-position.^[29]^ Not only *O*-MTs fall into this group, also *N*-MTs such as the anthranilate *N*-MT from *Ruta graveolens* (*Rg*ANMT, EC 2.1.1.111) clusters with class II enzymes (Figure 1).[^31^, ^32]^

**Figure 1.**
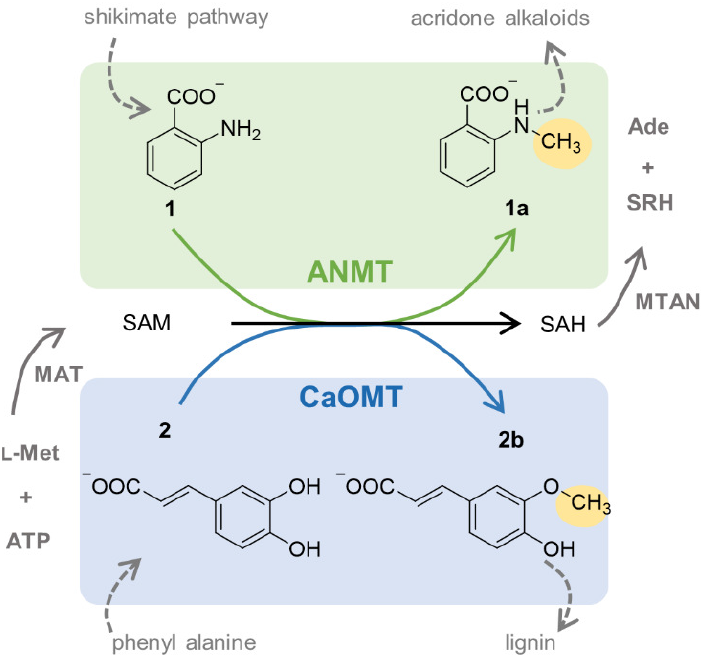
Three-enzyme cascade used in this work. L-methionine (L-Met) and ATP serve as starting material for the in situ synthesis of the cofactor SAM, catalysed by a MAT enzyme. The methyl group of SAM is transferred onto different substrates by an MT. The reaction catalysed by the ANMT *N-*methylates anthranilate (**1**) to *N-*methylantranilate (**1a)** (green). The natural reaction catalysed by the CaOMT is the methylation in 3-position of caffeate (**2**) to 3-methoxy-4-hydrocycinnamate **2b** (blue). In SAM-dependent methylation reactions the by-product SAH is formed, a known inhibitor for many MTs. Therefore, a third enzyme – MTAN – is used to cleave SAH into adenine and *S*-ribosyl-L-homocysteine (SRH) to shift the equilibrium to the product side.

The *N*-methylation of **1**, catalysed by the ANMT is a crucial step for the formation of acridone alkaloids and is important for plant growth.^[32]^ Despite the high similarity in amino acid sequences of the ANMTs and CaOMTs (>50% amino acid identity), the methyl acceptor atoms are different. Regarding the methylation mechanism, CaOMTs use a histidine residue as the catalytic base deprotonating the hydroxyl group that receives the methyl group of SAM.^[29]^ This histidine residue is found at an equivalent position in ANMTs.^[32]^ However, *Rg*ANMT has been described to exclusively catalyse the methylation of the amino group in **1**.^[32]^ In a recent paper describing an MT screening assay, initial insights into an extended substrate range towards other aniline substrates were reported.^[33]^

To our knowledge, only one ANMT has been described to date, which makes a bioinformatic analysis of the underlying molecular differences challenging. In this study, we analysed a panel of putative *N*- and *O*-selective MTs regarding their substrate scopes. The results led to the discovery of a new ANMT, and the detailed characterisation of aminophenols as substrates for ANMT as well as CaOMT enzymes.

## Results and Discussion

Based on homology searches for *Rg*ANMT using Protein BLAST, five enzymes with an amino acid identity > 50% were selected for use in further experiments (Table 1). Three candidates were from *Citrus sinensis* (*Cs*), one from *Prunus persica* (*Pp*), and one from *Medicago sativa* (*Ms*); the latter had already been characterised as a CaOMT (amino acid alignment in SI; Figure S 2).^[29]^

**Table 1.** Proteins selected from Protein BLAST using RgANMT as search sequence.

| Organism | Function | Identity [%]* | Similarity [%]* | Uniprot/ NCBI accession number | *N*-methylation | *O*-methylation |
| --- | --- | --- | --- | --- | --- | --- |
| *Ruta graveolens* (*Rg*) | ANMT<sup>[32]</sup> | 100 | 100 | A9X7L0.1 | +++ | - |
| *Citrus sinensis* (*Cs*) | ANMT | 81.4 | 88.8 | KAH9787224.1 | + | - |
| *Citrus sinensis* (*Cs*) |  | 75.5 | 82.3 | KDO86634.1 |  |  |
| *Prunus persica* (*Pp*) | CaOMT | 57.8 | 74.3 | XP_007218135.1 | - | +++ |
| *Citrus sinensis* (*Cs*) | CaOMT | 51.0 | 71.2 | XP_006494578.1 | - | ++ |
| *Medicago sativa* (*Ms*) | CaOMT<sup>[29]</sup> | 50.9 | 67.7 | P28002.1 | - | +++ |

\*referring to the amino acid sequence of *RgANMT*. The protein from *Citrus sinensis* (KDO86634.1) (grey) could not be used for activity assays due to solubility issues. +++ = full conversion (>99%); ++ = medium conversion (~50%); + = low conversion (<10%); - = no conversion

The cloning of *Rg*ANMT has been described previously.^[34]^ The *Escherichia coli* codon optimised synthetic genes encoding the other enzymes were cloned into pET-28a(+) vectors and were heterologously produced in *E. coli* BL21-Gold (DE3). All proteins contain an *N*-terminal His6-Tag and were purified *via* immobilised-metal affinity chromatography (IMAC). All proteins, except for one from *Citrus sinensis* (KDO86634.1) were soluble and tested for activity using **1** and **2** as substrates. All enzymes were active in the in vitro assays and identified as either an ANMT when accepting **1** or CaOMT when accepting **2** as a substrate (Table 1). Besides *Rg*ANMT, there was one other enzyme clearly identified as an ANMT, for which only **1** was a substrate. The other three enzymes accepted only **2** as a substrate, confirming their annotation as CaOMTs (Figure S 3).

### Substrate Scope

Comparing the results of the two ANMTs and three CaOMTs, *Rg*ANMT and *Pp*CaOMT showed the highest conversions for **1** and **2**, respectively. These two enzymes were then used in more detailed investigations with an extended substrate scope (Table 2). All compounds tested contained an amino- and hydroxyl group, making them potential substrates for both, *N*- and *O*-MTs. The third substituent was represented by a nitro (**3, 4**), bromo (**5, 6**), or chloro (**7, 8**) group either in the C-3 or C-4 positions.

**Table 2.**
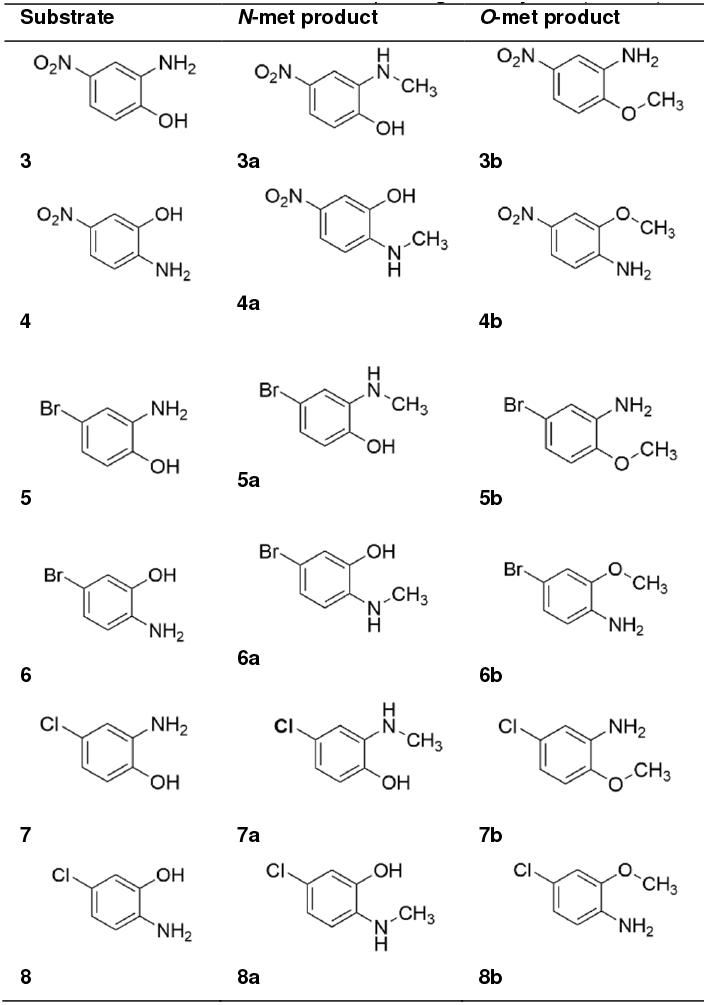
Substrates **3** – **8** and the corresponding *N-*methylated (**3a** – **8a**) and *O-*methylated (**3b** – **8b**) products described in this study.

| Substrate | *N*-met product | *O*-met product |
| --- | --- | --- |
| **3** | **3a** | **3b** |
| **4** | **4a** | **4b** |
| **5** | **5a** | **5b** |
| **6** | **6a** | **6b** |
| **7** | **7a** | **7b** |
| **8** | **8a** | **8b** |

*Pp*CaOMT accepted all substrates containing a phenolic group (Figure 2). The acceptance of substrates **3, 4** and **6** by *Rg*ANMT as suggested previously^[33]^ was confirmed; in addition, *Rg*ANMT also accepted substrates **5, 7**, and **8**. The corresponding HPLC traces show a high chemoselectivity of the two enzymes: the *N-* and *O-*alkylated products show shifts in retention time with the *N-*alkylated products eluting first (Figure 2 b; *Pp*CaOMT catalysed reaction). For the *O*-methylated products, the methylated compounds were available commercially, the *N*-methylated ones were produced enzymatically in vivo, purified and characterised with NMR spectroscopy (for details see SI). The chemoselectivity was additionally confirmed by ^13^C-NMR spectroscopy using ^13^C-labelled SAM produced from ^13^C-labelled L-methionine. The transfer of the ^13^C-labelled methyl group enables a distinct assignment of the the group acting as the nucleophile. In the NMR spectra of the *Rg*ANMT reaction, the signal at 29 ppm could be assigned to the carbon adjacent to the amino group. For the *Pp*CaOMT reaction, the new signal appeared at 56 ppm, confirming methylation in the hydroxyl position (Figure 2c; Figure S 12 – 15).

**Figure 2.**
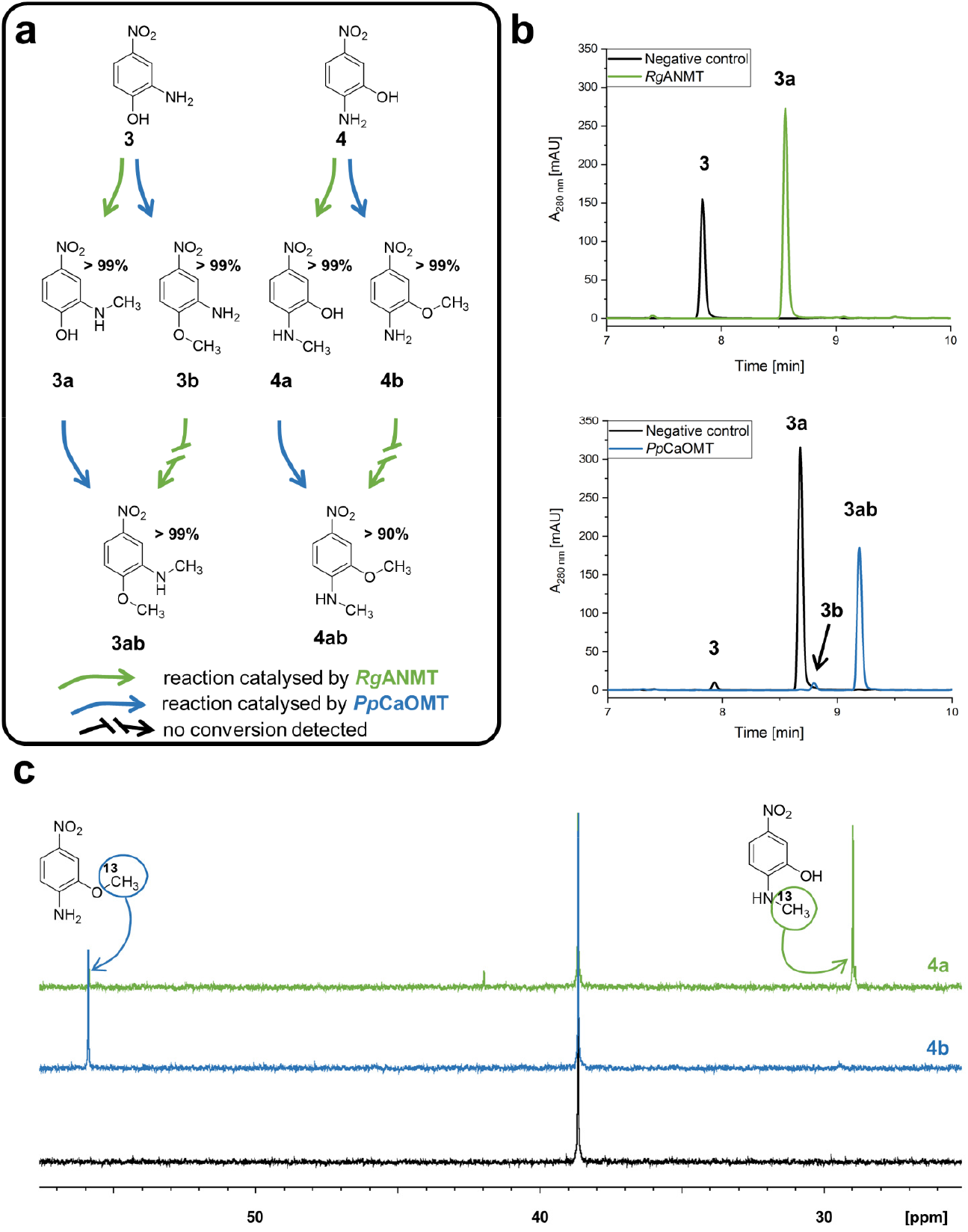
**a**: Formation of different products with conversion numbers starting with either substrates **3** and **4** using *Rg*ANMT (green) and *Pp*CaOMT (blue). **b**: Selected HPLC chromatograms representing the formation of **3a** catalysed by *Rg*ANMT starting with **3**, and the formation of **3ab** catalysed by *Pp*CaOMT starting with **3a** as substrate. **3a** was synthesised in vivo and purified before use. Small amounts of the starting material **3** were co-purified and methylated to product **3b** during the reaction. The reactions were stopped after 20 h. **c**: The chemoselectivity of the enzymes was confirmed by ^13^C-NMR experiments using ^13^C-labelled L-methionine in the enzyme cascade. The methyl group adjacent to the amino group gives a signal at 29 ppm and adjacent to the hydroxyl group at 56 ppm.

For both enzymes, full conversion of **3** and **4** into the single methylated products **3a/b** and **4a/b** was reached after 20 hours. The two substrates were used to record a time course for comparison of the enzyme velocities. The results showed that substrate **4** was converted more rapidly in both cases, with *Rg*ANMT being the faster enzyme converting **4** and **3** in 5 min and 40 min, respectively. For the *Pp*CaOMT reactions, full conversions occurred after 240 min for **4** and after 420 min for **3** (Figure 3; Figure S 6). The nitro substituents of both **3** and **4** have electron withdrawing effects; nevertheless, we did not recognise an impact on the biocatalytic methyl group transfer on either the amino or the hydroxyl group. In contrast to **3** and **4**, we did not observe full conversion for all of the halide containing substrates. After 20 h, amount of starting material **5** and **7** were still present, while **6** and **8** were fully converted to the products by both enzymes. Here, the positioning of the third substituent seems to be more important than for the nitro compounds, possibly due to interactions with several residues in the active site.

**Figure 3.**
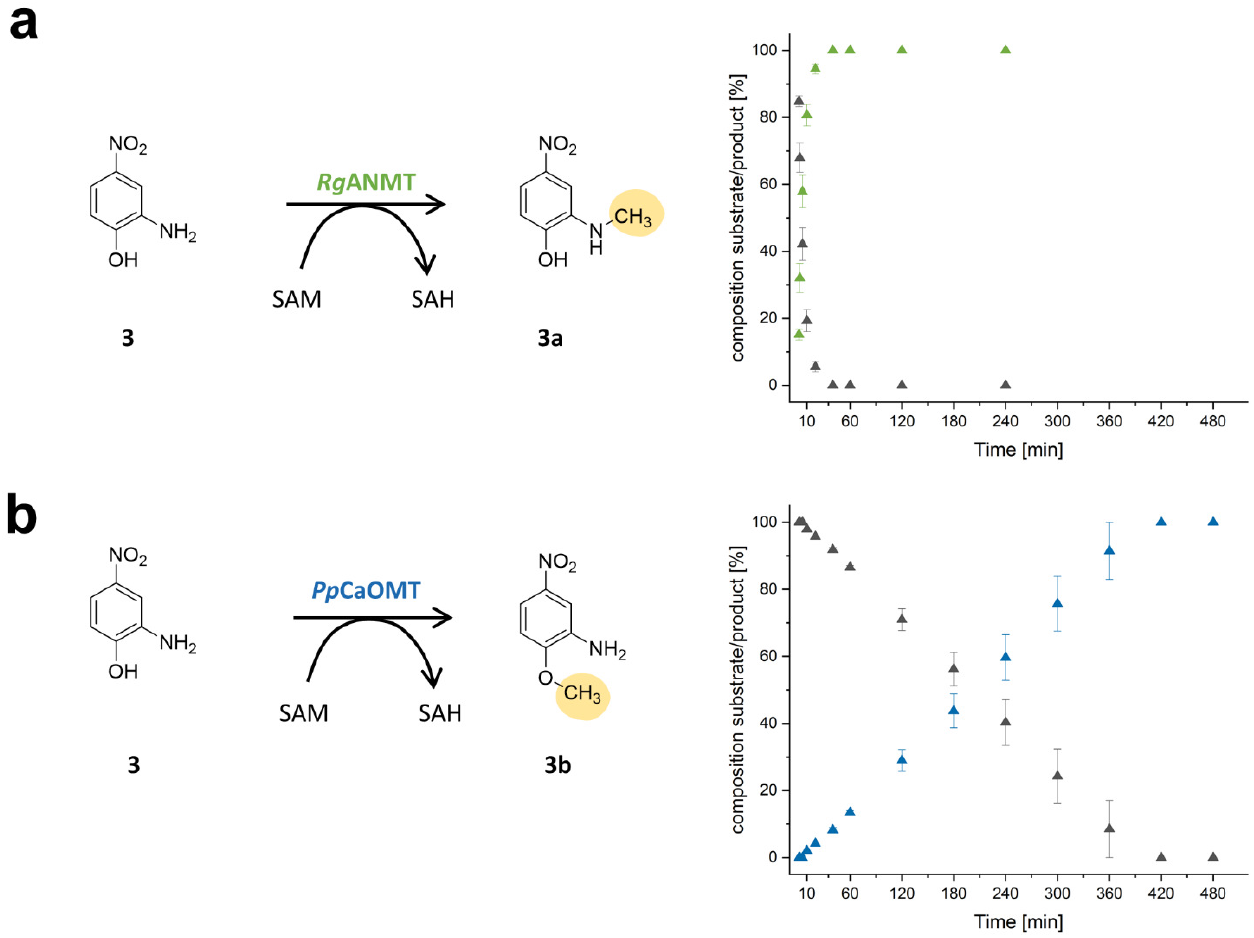
Timeline of methylation reaction for **3** to methylated products **3a** and **3b. a**: Reaction catalysed by *Rg*ANMT. Full conversion was reached after 40 min. The decrease of the substrate **3** (black) and increase of product **3a** (green) is shown after 1, 2, 5, 10, 20, 40, 60, 120 and 240 min. **b**: Reaction catalysed by *Pp*CaOMT. Full conversion was reached after 420 min. The decrease of the substrate **3** (black) and increase of product **3b** (blue) is shown after 1, 2, 5, 10, 20, 40, 60, 120, 240, 300, 360, 420 and 480 min.

In 2002, *Zubieta et al*. reported the crystal structure of *Ms*CaOMT. This enzymes has been described to accept a wide range of caffeic acid derivatives such as caffeoyl aldehyde and 5-hydroxyconiferaldehyde, methylating them at the 3-or 5-position.^[29]^ They proposed that the spacious active site is one reason for the acceptance of different molecules. This can be extended to the highly similar *Pp*CaOMT and *Rg*ANMT (Table 1) and might explain why the position of the hydroxyl and amino group is not decisive for substrate binding. The molecular reasons for the differing chemoselectivity are currently being explored in a follow-up study in our laboratory. Even though the two enzymes are similar regarding their amino acid sequences (and likely also their three-dimensional structures), the substrate positioning and or mechanism must differ leading to the high chemoselectivity of both enzymes.

Interestingly, when the *N-*methylated products **3a** and **4a** were incubated with the CaOMT they were accepted to achieve a second methylation step at the oxygen leading to full conversion for **3a** and conversions >90% for **4a** (Figure 2; Figure S 10 – S 11). In contrast to this, **3b** and **4b** were not accepted as substrates by *Rg*ANMT. This might be due to differences in *Rg*ANMT’s active site hindering access by the bulkier *O*-methylated compounds or correct binding mode near the cofactor.

### Preparative Synthesis of Selected Compounds

The *N*-methylated products for all used substrates are not commercially available. We therefore decided to elucidate the potential of *Rg*ANMT as a catalyst for the preparative enzymatic synthesis of **3a** and **4a**. An in vitro and an in vivo approach was set up for comparison purposes. The in vitro experiments were performed at a 20 mL scale using 10% (V/V) crude clarified lysate of all enzymes involved in the three-enzyme cascade. ATP and L-methionine were added in excess (0.2 mmol), and 0.1 mmol of **3** or **4** was used as the methyl acceptor. After 20 h of incubation at 37 °C the reaction was stopped and the products were extracted and purified *via* preparative HPLC. The purified yield of the products **3a** and **4a** were 62% and 34%, respectively. For the in vivo approach, **3** and **4** were methylated with *E. coli* BL21-Gold(DE3) whole cells transformed with pET28a(+)::*rg*anmt. A culture with an OD600 of 3 in MM9-medium was incubated with 4 mmol L-methionine and 0.15 mmol of **3** or **4** for 48 h at 30 °C followed by preparative purification *via* Puriflash. In the in vivo experiment the yield for **3a** and **4a** were 43% and 19%, respectively. NMR spectroscopic analysis confirmed the formation of the *N*-methylated products for both experiments (Figure S 16 – S 19). In the HPLC traces, small amounts of the starting material remained using both methods (Figure S 9). Optimised purification steps might increase the yields and purity for all experiments. Both methods were successful regarding the production of the desired substances (Table 3). In the in vitro approach, the required compounds can be added more precisely; and this also presents a straight-forward opportunity to use cofactor analogues as described before.^[35]^ In the in vivo experiment, the cofactor ATP is provided by the cell. Also, cells can be kept alive and be used for a continuous process while enzymes used in in vitro studies will lose activity over time. Substrates unable to pass the bacterial cell wall will not be methylated, as the cofactor and enzyme are only present inside of the cell.

**Table 3.** Comparison of in vitro and in vivo upscale experiments.

|  | in vitro experiment | in vivo experiment |
| --- | --- | --- |
| Reaction volume | 20 mL | 200 mL |
| ATP | 0.2 mmol | - |
| L-methionine | 0.2 mmol | 4 mmol |
| Substrate (**3** / **4**) | 0.1 mmol | 0.15 mmol |
| Purified yield | **3a**: 62% | **3a**: 43% |
|  | **4a**: 34% | **4a**: 19% |
| Purity | **3a**: 96% | **3a**: 97% |
|  | **4a**: 90% | **4a**: 85% |

### Alkylation Reactions

Besides methylation, the three-enzymes cascade has already been used for the transfer of bulkier alkyl chains in previous work.^[19–22]^ For the in situ synthesis of SAM derivatives, we used L-ethionine or *S*-allyl-L-homocysteine together with ATP [forming either *S*-adenosyl-L-ethionine (SAE) as ethyl group donor, or *S*-adenosyl-*S*-allyl-L-homocysteine (SAA) as allyl group donor]. Instead of *Ec*MAT, the MAT from *Thermococcus kodakarensis* (*Tk*) was employed since previous data suggested its superior suitability for the transfer of bulkier alkyl chains.^[19,36]^ In earlier work, it was shown that *Rg*ANMT accepts modified cofactors and can transfer ethyl groups onto **1**.^[14]^

Experiments were performed to determine if *Rg*ANMT and *Pp*CaOMT can use SAE and SAA to transfer the ethyl-or allyl group onto the unnatural substrates **3** and **4**. Along with consumption of the substrate, the HPLC traces showed a growing peak corresponding to adenine, and a new peak increasing in intensity, which was assigned to the ethylated and allylated products by LC-MS analysis, confirming successful alkylation reactions (Figure 4, Figure S 5; S 7). *Rg*ANMT preferably *N-*ethylated **3** to **3c** with conversions of 41% (+/-2%) and *Pp*CaOMT *O-*ethylated **4** to give **4d** in up to 36% (+/-2%) (analysis by HPLC) (Figure S 8). Conversions for the other products formed were between 15 – 30 % supporting that both *Rg*ANMT and *Pp*CaOMT can use SAE and SAA to transfer the ethyl-or allyl group also onto the unnatural substrates **3** and **4**, thereby increasing the pool of products accessible by the formation of compounds **3c-f** and **4c-f** (Figure 4).

**Figure 4.**
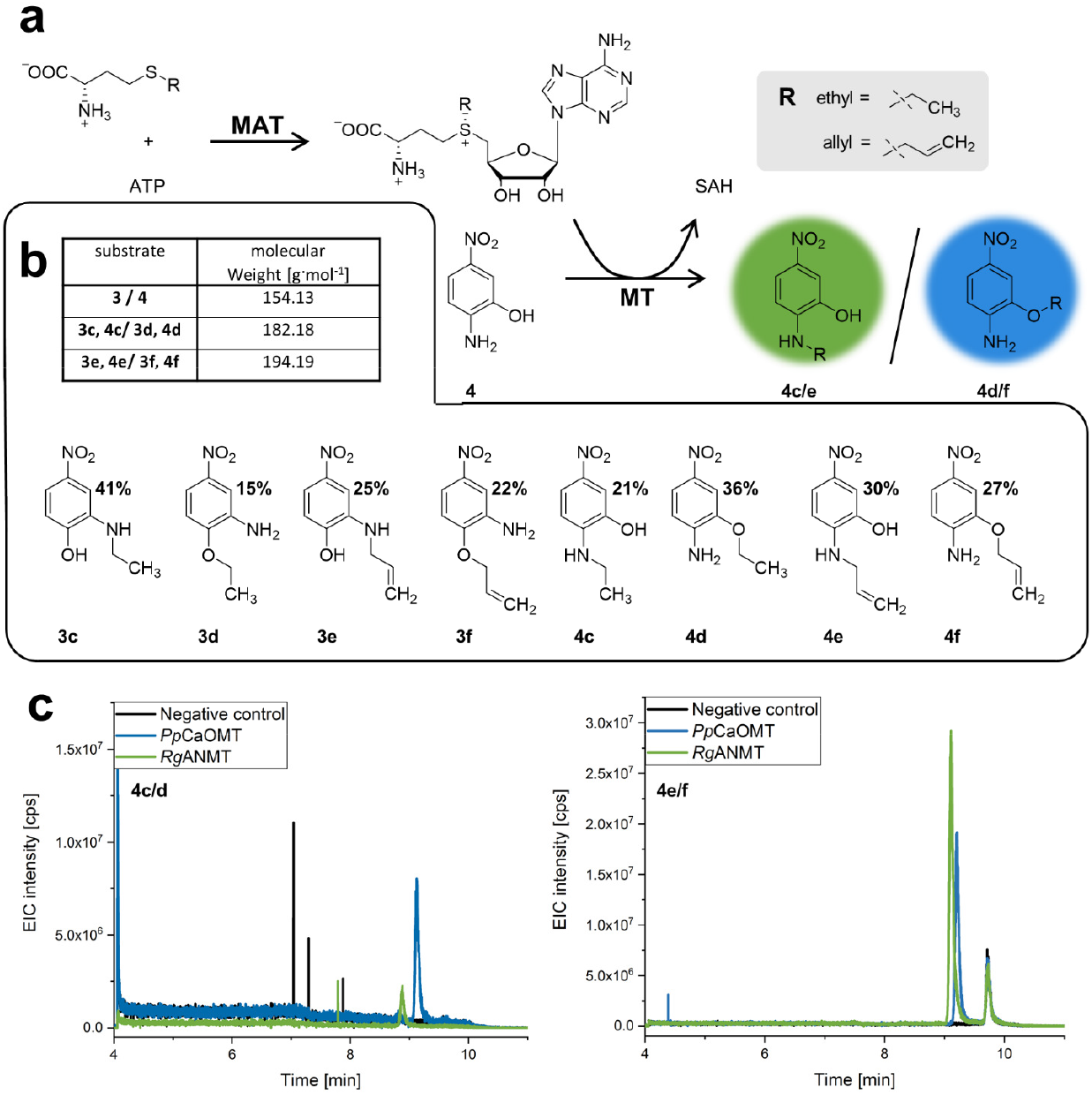
Generation of SAM derivatives to transfer bulkier alkyl chains by different MTs onto substrate **a**: ATP and L-ethionine (**Re**) or L-allylhomocysteine (**Ra**) were used for the cofactor generation. The cofactor derivative is then used by the MT to transfer the ethyl or allyl group onto the substrate. **b**: Products with ethyl and allyl groups (**3c-3f** and **4c-4f**) with conversion yields calculated by HPLC are shown. **c**: In the extracted ion chromatogram the mass of the formed ethylated [**4c** and **4d** (183.18 Da in positive mode)] and allylated products **4e** and **4f** (195.19 Da in positive mode) catalysed by *Rg*ANMT (green) and by *Pp*CaOMT (blue) are displayed.

## Conclusion

Here, we described a set of novel enzymes related to caffeic acid *O*-MTs, including a new ANMT from *C. sinensis*. Notably, *Pp*CaOMT and *Rg*ANMT were established as promiscuous biocatalysts for small molecule methylation. Besides their natural substrates, both enzymes accept a broad range of non-physiological substrates and catalyse their methylation chemoselectively in the presence of different nucleophilic species. Although both enzymes show high similarities in their amino acid sequence, both reactions interestingly lead to single *O*-or *N-*methylated products. *Pp*CaOMT accepted the produced *N*-methylated products **3a** and **4a** forming the double methylated products **3ab** and **4ab**. A closer investigation of the active sites by co-crystallisation and computational methods as well as mutagenesis experiments will be carried out in the future with the aim to understand the differences in the enzymes’ mechanisms leading to highly pure *N-* and *O-*alkylated products.

In addition to SAM, both enzymes accept analogues of the natural cofactor, such as the ethyl- and allyl-SAM. Other L-methionine derivatives can be used in the future to expand the product pool. Recently, a four enzyme cascade was published, starting from thiol compounds to produce different L-methionine derivatives and subsequent transfers catalysed by MTs.^[37]^ Adapting this enzyme cascade will increase the pool of products even further. Finally, the methylation reactions were performed on a larger scale in vitro, and compared to an in vivo strategy using whole cells. The yield of the isolated products was between 19 – 62%. Both methods led to the desired products, with none of them having clear advantages regarding yields. The decision of which method to use will therefore depend on the individual case, for example, using compounds that do not enter the bacterial cells. Also, the use of SAM analogues will require the in vitro method; while the in vivo option might be better suited for a cost-efficient synthesis of methylated products, as no addition of a cofactor building block is required.

## Acknowledgements

The work of the Andexer group was supported by the Deutsche Forschungsgemeinschaft (RTG1976 for D.P. and E.J. and Heisenberg program for J.N.A.), the European Research Council in frame of the Horizon 2020 program (ERC starting grant 716966) and the Deutsche Bundesstiftung Umwelt (DBU PhD scholarship for M.K.F.M). Work in the Andexer and Hailes groups was supported by the EU (BMBF grant 161B0626B; BBSRC grant BB/R021643/1 for F.S.) through the ERACo Biotech project BioDiMet. E.M.C. was funded by the Engineering and Physical Sciences Research Council (EPSRC), grant EP/N509577/1. We thank the whole BioDiMet team for helpful discussions in the consortium. From the University of Freiburg, Sascha Ferlaino is acknowledged for help with NMR analysis, and Adelheid Nagel, Kalle Kind, Aaliya Afandi Lee and Katharina Strack for technical assistance with protein production and time course experiments. Dr Dipali Mhaindarkar is acknowledged for initial work on in vivo preparation of methylated standards, as well as Prof. Michael Müller for useful advice and comments regarding enzyme selectivity. Additionally, the EPSRC for 700 MHz NMR equipment support (EP/P020410/1) at UCL.

## Additional Information

### Supplementary information

is available online.

