## Supplemental Material for "Expanding the Substrate Scope of *N*- and *O*-Methyltransferases from Plants for Chemoselective Alkylation"

### Experimental section

#### Materials

All substrates and reference compounds were purchased from Sigma-Aldrich (ATP, SAM, SAH, L-methionine, L-methionine-(methyl-$^{13}$C), L-ethionine, 2-amino-4-bromo phenol, 2-amino-5-bromo phenol, 2-amino-4-chloro phenol, 2-amino-5-chloro phenol, 2-amino-4-nitro phenol, 2-amino-5-nitro phenol, 4-chloro-2-methoxyaniline, 5-chloro-2-methoxyaniline, 2-methoxy-4-nitro aniline, 2-methoxy-5-nitro aniline, caffeic acid, ferulic acid, isoferulic acid, 3,4-dihydroxybenzoic acid), Alfa Aesar (*N*-methylanthranilic acid) and Fluka (anthranilic acid) in the highest purity available. Buffer ingredients, as well as cultivation media were obtained by Carl Roth.

**Table S 1** – Used substrates and products with abbreviations and retention times. A, B, and C refers to the different analytical methods.

| **Substrate** | **Abbreviation / #** | **Retention time [min]** | | |
|---|---|---|---|---|
| | | **A** | **B** | **C** |
| L-methionine | **L-met** | - | - | - |
| L-ethionine | **L-eth** | - | - | - |
| *S*-allyl-L-homocysteine | **L-all** | - | - | - |
| adenosine triphosphate | **ATP** | 1.9 | 1.1 | - |
| *S*-adenosyl-L-methionine | **SAM** | 1.7 | 1.3 | - |
| *S*-adenosyl-L-homocysteine | **SAH** | 8.1 | 1.2 | - |
| adenine | **ade** | 9.3 | 1.6 | - |
| *S*-adenosyl-L-ethionine | **SAE** | - | 1.3 | - |
| adenosyl-*S*-allyl-L-homocysteine | **SAA** | - | 1.4 | - |
| anthranilic acid | **1** | 3.6 | - | - |
| *N*-methyl anthranilic acid | **1a** | 6.8 | - | - |
| caffeic acid | **2** | 3.2 | - | - |
| 3-methoxy-4-hydroxycinnamic acid | **2b** | 6.4 | - | - |
| 3-hydroxy-4-methoxycinnamic acid | **2b''** | 7.8 | - | - |
| 2-amino-4-nitrophenol | **3** | - | 7.8 | 5.0 |
| 2-(methylamino)-4-nitrophenol | **3a** | - | 8.6 | - |
| 2-methoxy-5-nitroanilin | **3b** | - | 8.7 | 6.3 |
| 2-methoxy-*N*-methyl-5-nitroaniline | **3ab** | | 9.2 | - |
| 2-(ethylamino)-4-nitrophenol | **3c** | - | 8.8 | - |
| 2-ethoxy-5-nitroaniline | **3d** | - | 9.0 | - |
| 2-(allylamino)-4-nitrophenol | **3e** | - | 8.9 | - |
| 2-allyloxy-5-nitroaniline | **3f** | - | 9.1 | - |
| 2-amino-5-nitrophenol | **4** | - | 8.2 | 6.0 |
| 2-(methylamino)-5-nitrophenol | **4a** | - | 8.7 | - |
| 2-methoxy-4-nitroanilin | **4b** | - | 8.8 | 6.7 |
| 2-methoxy-*N*-methyl-4-nitroaniline | **4ab** | | 9.1 | - |
| 2-(ethylamino)-5-nitrophenol | **4c** | - | 8.9 | - |
| 2-ethoxy-4-nitroanilin | **4d** | - | 9.0 | - |
| 2-(allylamino)-5-nitrophenol | **4e** | - | 8.9 | - |
| 2-allyloxy-4-nitroanilin | **4f** | - | 9.1 | - |
| 2-amino-4-bromophenol | **5** | - | 7.9 | - |
| 4-bromo-2-(methylamino)phenol | **5a** | - | 8.7 | - |
| 5-bromo-2-methoxyaniline | **5b** | - | 9.0 | - |
| 2-amino-5-bromophenol | **6** | - | 7.1 | - |

| Substrate | Abbreviation / # | Retention time [min] | | |
|---|---|---|---|---|
| | | A | B | C |
| 5-bromo-2-(methylamino)phenol | **6a** | - | 7.9 | - |
| 4-bromo-2-methoxyaniline | **6b** | - | 8.8 | - |
| 2-amino-4-chlorophenol | **7** | - | 7.3 | - |
| 4-chloro-2-(methylamino)phenol | **7a** | - | 8.5 | - |
| 5-bromo-2-methoxyaniline | **7b** | - | 8.9 | - |
| 2-amino-5-chlorophenol | **8** | - | 6.0 | - |
| 5-chloro-2-(methylamino)phenol | **8a** | - | 7.4 | - |
| 4-bromo-2-methoxyaniline | **8b** | - | 8.5 | - |

##### Synthesis of *S*-allyl-L-homocysteine

*S*-allyl-L-homocysteine was used for the in situ synthesis of the cofactor derivative adenosyl-*S*-allyl-L-homocysteine (SAA). Since this compound is not commercially available, the synthesis was performed according to previous work.[1] The purity was confirmed by $^1$H-NMR spectroscopic analysis (**Figure S 20**).

#### Methods

##### Cloning

The *Ec*MAT, *Tk*MAT, *Rg*ANMT, and *Ec*MTAN plasmids have been described in previous publications.[2,3]

The synthetic genes encoding for the other enzymes were purchased from Invitrogen (Thermo Fisher Scientific, Waltham, MA, USA). The primers used for amplification were ordered at Eurofins (**Table S 2**). After amplification, PCR samples were analysed by agarose gel electrophoresis (1% agarose, 100 V, 60 min). For the cloning procedure an In-Fusion protocol from Takara Bio Europe (Saint-Germain-en-Laye, France) was used. For In-Fusion cloning, restriction sites are not necessary. Still, there were restriction sites added.

**Table S 2** – Primers for gene amplification and linearisation of pET28a(+).

| Primer | Sequence 5'-3' |
|---|---|
| KAH9787224.1 (*Citrus sinensis*)_fw | CGCGCGGCAGC<u>CATATG</u>GGTAGCCTGAGCGAATATCA |
| KAH9787224.1 (*Citrus sinensis*)_rev | GTGCGGCCGC<u>AAGCTT</u>ATTTGAAAAATTCCATGATATACA |
| KDO86634.1 (*Citrus sinensis*)_fw | CGCGCGGCAGC<u>CATATG</u>GGTAGCCTGAGCGAATA |
| KDO86634.1 (*Citrus sinensis*)_rev | GTGCGGCCGCAAGCT<u>AAGCTT</u>ATTTGAAAAATTCCATGAT |
| XP_007218135.1 (*Prunus persica*)_fw | CGCGCGGCAGC<u>CATATG</u>GCAAGCAGCCTGGAAC |
| XP_007218135.1 (*Prunus persica*)_rev | GTGCGGCCGCAAGCT<u>AAGCTT</u>ATTTGAAAAATTCCATCAC |
| XP_006494578.1 (*Citrus sinensis*)_fw | CGCGCGGCAGC<u>CATATG</u>GATAGCATTGTTGATGGTG |
| XP_006494578.1 (*Citrus sinensis*)_rev | GTGCGGCCGCAAGCT<u>AAGCTT</u>ATTTGTAAAATTCCATAAC |

| | |
|---|---|
| P28002.1 (*Medicago sativa*) fw | CGCGCGGCAGCCATATGGGTAGCACCGGTGAAAC |
| P28002.1 (*Medicago sativa*) rev | GTGCGGCCGCAAGCTAAGCTTACACTTTTTAAGGAATTC |
| *pET28a_Nde*I | TATGGCTGCCGCGCGGCACC |
| *pET28a_Hind*III | AGCTTGCGGCCGCACTCGAG |

*Restriction sites are underlined.

For the PCR, 20 ng of the DNA template (gene or vector) was used. Additionally, the samples contained 2.5 µL of forward and reverse primer (10 µmol · $L^{-1}$), 25 µL 2X Phusion Flash PCR Master Mix (Thermo Fisher), and were filled up to 50 µL with $H_2O_{millipore}$. A 3-step PCR protocol was performed (**Table S *3***).

**Table S 3** – 3-step PCR protocol.

| Step | Temperature [°C] | Time [s] | Cycles |
|---|---|---|---|
| Initial denaturation | 98 | 30 | 1 |
| Denaturation | 98 | 10 | 30 |
| Annealing | 55 | 30 | |
| Elongation | 72 | 90 | |
| Final elongation | 72 | 240 | 1 |
| Storage | 8 | ∞ | |

For the In-Fusion cloning, 50 ng of the linearised vector and 100 ng of the amplified gene were combined with 2 µL of 5X In-Fusion HD Enzyme Premix, and filled up to 10 µL with $H_2O_{millipore.}$ The reaction took place at 50 °C for 15 min. Afterwards, a standard chemical transformation into *E. coli* Stellar cells was performed. Successful cloning was confirmed by Sanger Sequencing (Eurofins Genomics).

#### Protein overproduction

The plasmids were transformed into E. *coli* BL21-Gold(DE3) cells. One colony was incubated in 5 mL LB-medium containing kanamycin (50 µg · $L^{-1}$) at 37 °C, 170 rpm overnight. The pre culture (1%) was added to 400 mL LB-medium containing kanamycin (50 µg · $L^{-1}$) and incubated at 37 °C, 170 rpm until an $OD_{600}$ of 0.5 – 0.7 was reached. Isopropyl-β-D-thiogalactopyranoside (IPTG) was added to a final concentration of 0.25 mM for overexpression induction. The overproduction took place at 20 °C, 140 rpm for 20 h. Cells were harvested by centrifugation (4 °C, 8000 rpm, 7.8 x 1000 g, 20 min) and stored at 4 °C until following purification.

For test expression experiments cell lysis was performed using BugBuster (Merck AG) according to the manufacturer's protocol. Proteins were visualised *via* SDS-PAGE analysis.

#### Protein purification

Pellets were resuspended (4 mL · $g^{-1}$) in lysis buffer [40 mM Tris-HCl, pH 8.0, 100 mM, NaCl, 10% (w/v) glycerol]. Cell lysis was performed *via* sonication (Branson Sonifier 250, Emerson,

St. Louis, MO, USA [duty cycle 50%, intensity 50%, 5 x 30 s with 30 s breaks within]). The lysis solution was centrifuged for 40 min at 4 °C (24.9 x 1000 g) to precipitate non-soluble cell fragments. The crude lysate was applied to a nickel-NTA column followed by washing and elution steps. First, 5 mL of lysis buffer was used containing 5, 10, 20, 50 mM imidazole, followed by elution steps with 5 mL of lysis buffer containing 100, 150, 200, 300 mM imidazole. Later, it was shown that a reduced two-step purification was sufficient. Here, 30 mL lysis buffer containing 10 mM imidazole were used for washing, followed by the elution step using 20 mL lysis buffer with 250 mM imidazole. After, the protein solution was desalted, using PD-10 columns (GE Healthcare Life Sciences, Little Chalfont, UK) according to the manufacturer's instructions. Protein concentration was determined with a NanoDrop 2000 (Thermo Fisher Scientific, Waltham, MA, USA) at 280 nm. The molecular weight and extinction coefficient (including $His_6$-tag) was calculated with the ExPASy ProtParam tool for further concentration measurements *via* nanodrop.[4]

#### Assays

In general, the qualitative and quantitative assays were performed at least in triplicates. The in vivo and in vitro syntheses were performed one time and confirmed by NMR analysis.

in vitro assays (small scale)

A standard assay for the characterisation of the MTs and the substrate screening was prepared in 200 – 500 µL with 50 mM Tris pH 7.5, 20 mM $MgCl_2$, and 50 mM KCl. 10 µM MAT and MT and 2 µM MTAN enzymes were used. 3 mM ATP and L-methionine (or L-ethionine or L-allylhomocysteine) and 2 mM substrate was added. Assays were incubated at 37 °C, 300 rpm and samples were taken after 1 and 20 h. The reaction was stopped by the addition of perchloric acid (total conc. 2.5%) and the samples stored at -4 °C until analysis. For samples analysed by $^{13}$C-NMR, $^{13}$C-labelled L-methionine was used as substrate.

For the time course assay, the enzyme and substrate concentrations were adapted due to solubility issues of the *O*-methylated compounds. The concentration of the MAT and MT enzyme was 3 µM and of MTAN 1 µM. ATP and L-methionine concentration was 1 mM and MT substrate concentration was 0.5 mM.

Conversion rates of the quantitative assays were calculated from substrate and product AUC at certain time points.

$$Conversion\ \% = \frac{AUC(product)}{AUC\ (substrate) + AUC\ (product)} * 100$$

in vitro assays (upscale)

The reaction volume of the in vitro upscale experiments was 20 mL. 2 mL of a stock solution [100 mM ATP and L-methionine in 50 mM HEPES (pH 7.5)] was added to substrate **3** or **4** (0.1 mmol, 15.4 mg). 10% (v/v) of *Ec*MAT and *Rg*ANMT and 2% (v/v) of *Ec*MTAN lysate was added. The solution also contained 20 mM $MgCl_2$ and 200 mM KCl. The reaction was incubated at 37 °C and 180 rpm overnight. After centrifugation, the supernatant was extracted with ethyl acetate (3x 20 mL). The combined organic fractions were evaporated under vacuum. The residue was dissolved in a water acetonitrile mixture (9:1) and purified via HPLC method D . Product fractions were freeze-dried.

in vivo assays (upscale)

For the in vivo synthesis of the *N*-methylated products **3a** and **4a** pre and main cultures with E*. coli* BL21-Gold(DE3) cells containing pET28a(+)::*rganmt* were prepared as described for protein overproduction. Instead of 0.25 mM, 1 mM IPTG was used for the induction of overexpression. For cell harvest, the main cultures were incubated on ice for 10 min and centrifuged for 20 min at 4 °C (2500 x g). After discarding the supernatant, the cell pellets were resuspended in precooled MM9 medium to an $OD_{600}$ of 3.0. IPTG (1 mM) and kanamycin (50 µg · $mL^{-1}$) were added. 200 mL of the cultures were transferred in 500 mL shaking flasks with baffles. For the following in vivo reaction sterile filtered L-methionine (20 mM final concentration) was added. 750 µM of the MT substrate was added (stock solution 100 mM in DMSO). The reaction took place at 37 °C, 170 rpm for 48 h. The product was extracted using ethyl acetate (3x equal volume). Afterwards, the organic phase was separated from the cells and evaporated in a rotary evaporator. Purification of the *N*-methylated products was performed with a PuriFlash system PF_XS_520 and a Biotage SNAP Cartridge (KP-Sil, 50g). A flow of 15 ml/min was used and the gradient was performed with cyclohexane (Solution A) and ethyl acetate (Solution B) starting with 80% A until 5 min, lowering to 70% A until 7 min, 60% A until 9 min, 50% A until 11 min and holding until 14.2 min, then lowering to 20% A until 16 min and holding until 17 min, lowering to 0% A until 17 min and holding until 20 min. The formation of the products was confirmed by NMR analysis (**Figure S 16** – **Figure S 17**).

#### HPLC analysis

Four different methods were used for HPLC analysis. **Method A** was used for the characterisation of the different MT candidates methylating either substrate **1** or **2**. This method has been described before.[3] **Method B** was used for the analysis of the substrate screen and the in vivo experiments and has been described recently.[5] The scaled up reactions were analysed and purified using **Method C** and **D** (**Table S4**)

**Table S 4** – Methods used for upscale reaction analysis and purification

| | **Method C** | **Method D** |
|---|---|---|
| Assay | Analytical method of Up scale reaction | Purification method of Up scale reaction |
| HPLC system | Agilent 1260 infinity II | Agilent 1260 InfinityTM |
| Column | ACE 5-C18 300 column (4 µm, 4.6 mm × 150 mm) | SupelcoTM Discovery BIO wide pore (C18, 10 µm, 2.12 cm x 25 cm) |
| Mobile phase A | 0.1% TFA | 0.1% TFA |
| Mobile phase B | acetonitrile | acetonitrile with 0.1% TFA |
| Detection wavelenght | 280 nm | 280 nm |
| Flow rate | 1 mL · $min^{-1}$ | 8 mL · $min^{-1}$ |
| Injection volume | 10 µL | 500 – 900 µL |
| Gradient | 1 min 90% A/ 10% B<br>6 min to 15% A/ 85% B<br>0.1 min to 0% A/ 100% B<br>1.9 min 0% A/ 100% B<br>0.1 min to 90% A/ 10% B<br>2.4 min 90% A/ 10%B | 3 min 90% A/ 10% B<br>7 min to 50% A/ 50% B<br>15 min to 0% A/ 100% B<br>5 min 0% A/ 100% B<br>1 min 90% A/ 10% B<br>4 min 90% A/ 10% B |

#### LC-MS analysis

An LC-MS method that has been described earlier to analyse MT substrates and products was used to analyse the ethylated and allylated products of the MT reactions.[5] The samples were diluted 1:50 and filtered before measurement. A Q1 scan was performed searching for the masses of the mother ion fragments (**Table S 5**) in positive mode.

**Table S 5** – Investigated mother fragments in LC-MS analysis for confirmation of ethylation and allylation.

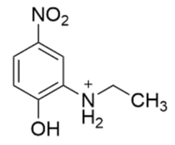

**3c**

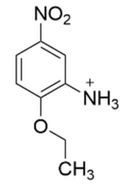

**3d**

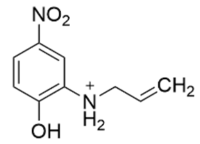

**3e**

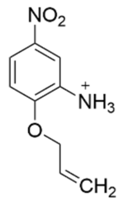

**3f**

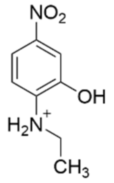

**4c**

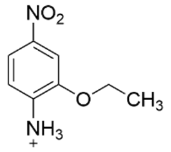

**4d**

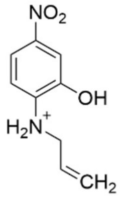

**4e**

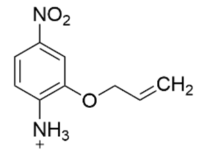

**4f**

*m/z of $[M+H]^+$ of products **3c/d** and **4c/d**: 183.18 Da and of products 3e/f and 4e/f: 195.19 Da

#### NMR analysis

A Bruker Avance III HD 400 MHz instrument was used to analyse the chemoselective substrate methylation and the in vivo synthesis of the *N*-methylated amino nitrophenols by $^{13}C$-NMR analysis. The upscaled reactions and synthesis of *S*-allyl-L-homocysteine were measured by $^{13}C$- and $^{1}H$-NMR analysis using Bruker Avance Neo 500 and Bruker Avance Neo 700 spectrometers.

In **Table S 6** the $^{13}C$ signals for the used assay compounds are listed. The signals for *N*- and *O*-methylation are displayed in each NMR figure (**Figure S 10 – 15**).

**Table S 5** – $^{13}C$-NMR signals for compounds used in the methylation assays.

| | $\delta$ [ppm] |
|---|---|
| Methionine (S-$\underline{C}H_3$) | 13.8 |
| SAM (S-$\underline{C}H_3$) | 23.5 |
| Methionine ($H_2N$-$\underline{C}H$) | 59.4 |
| Tris (HO-$\underline{C}H_2$) | 61.6 |
| Glycerol ($\underline{C}H_2$) | 62.7 |
| Glycerol ($\underline{C}H$) | 72.1 |

#### Protein sequences

The following protein sequences and enzyme sizes include the His$_6$-tag (marked in bold).

>*Ec*MAT (44.1 kDa)

MGSS**HHHHHH**SSGLVPRGSHMAKHLFTSESVSEGHPDKIADQISDAVLDAILEQDPKARVACETYVKT
GMVLVGGEITTSAWVDIEEITRNTVREIGYVHSDMGFDANSCAVLSAIGKQSPDINQGVDRADPLEQG
AGDQGLMFGYATNETDVLMPAPITYAHRLVQRQAEVRKNGTLPWLRPDAKSQVTFQYDDGKIVGIDAV
VLSTQHSEEIDQKSLQEAVMEEIIKPILPAEWLTSATKFFINPTGRFVIGGPMGDCGLTGRKIIVDTY
GGMARHGGGAFSGKDPSKVDRSAAYAARYVAKNIVAAGLADRCEIQVSYAIGVAEPTSIMVETFGTEK
VPSEQLTLLVREFFDLRPYGLIQMLDLLHPIYKETAAYGHFGREHFPWEKTDKAQLLRDAAGLK

>*Tk*MAT (46.7 kDa)

MGSS**HHHHHH**SSGLVPRGSHMAGKVRNIVVEELVRTPVEMQKVELVERKGIGHPDSIADGIAEAVSRA
LSREYVKRYGIILHHNTDQVEVVGGRAYPQFGGGEVIKPIYILLSGRAVEMVDREFFPVHEIALKAAK
DYLRKAVRHLDLEHHVIIDSRIGQGSVDLVGVFNKAKKNPIPLANDTSFGVGYAPLSETEKIVLETEK
YLNSDEFKKKYPAVGEDIKVMGLRKGDEIDLTIAAAIVDSEVDNPDDYMAVKEAIYEAAKGIVESHTE
RPTNIYVNTADDPKEGIYYITVTGTSAEAGDDGSVGRGNRVNGLITPNRHMSMEAAAGKNPVSHVGKI
YNILSMLIANDIAEQVEGVEEVYVRILSQIGKPIDEPLVASVQIIPKKGYSIDVLQKPAYEIADEWLA
NITKIQKMILEDKVNVF

>*Ec*MTAN (26.5 kDa)

MGSS**HHHHHH**SSGLVPRGSHMKIGIIGAMEEEVTLLRDKIENRQTISLGGCEIYTGQLNGTEVALLKS
GIGKVAAALGATLLLEHCKPDVIINTGSAGGLAPTLKVGDIVVSDEARYHDADVTAFGYEYGQLPGCP
AGFKADDKLIAAAEACIAELNLNAVRGLIVSGDAFINGSVGLAKIRHNFPQAIAVEMEATAIAHVCHN
FNVPFVVVRAISDVADQQSHLSFDEFLAVAAKQSSLMVESLVQKLAHG

>*Pp*CaOMT (43.7 kDa)

MGSS**HHHHHH**SSGLVPRGSHMASSLERKSHPKINHAEPEDEITKEEEDESFCYAMQLVGSSVLSMSLQ
SAIKLGIFDIIARKGPGAKLSSSEIATKIGTENPEAPVMVDRILRLLTSHSVLNCSAVAANGGSDFQR
VYSLGPVSKYFVNDEEGGSLGPLLTLIQDRVFLESWSQLKDAVVEGGIPFNRVHGMHAFEYPGLDPRF
NQVFNTAMFNHTTIVIKKLLHIYKGLEDKNLTQLVDVGGGLGVTLNLITSRYQHIKGINFDLPHVVNH
APSYPGVEHVGGDMFASVPSGDAIFMKWILHDWSDEHCLKLLKNCYKAIPDNGKVIVVEALLPAMPET
STATKTTSQLDVLMMTQNPGGKERSEQEFMALATGAGFSGIRYECFVCNFWVMEFFK

>*Rg*ANMT (42.2 kDa)

MGSS**HHHHHH**SSGLVPRGSHMGSLSESHTQYKHGVEVEEDEEESYSRAMQLSMAIVLPMATQSAIQLG
VFEIIAKAPGGRLSASEIATILQAQNPKAPVMLDRMLRLLVSHRVLDCSVSGPAGERLYGLTSVSKYF
VPDQDGASLGNFMALPLDKVFMESWMGVKGAVMEGGIPFNRVHGMHIFEYASSNSKFSDTYHRAMFNH
STIALKRILEHYKGFENVTKLVDVGGGLGVTLSMIASKYPHIQAINFDLPHVVQDAASYPGVEHVGGN
MFESVPEGDAILMKWILHCWDDEQCLRILKNCYKATPENGKVIVMNSVVPETPEVSSSARETSLLDVL
LMTRDGGGRERTQKEFTELAIGAGFKGINFACCVCNLHIMEFFK

>*Cs*ANMT (41.9 kDa)

MGSS**HHHHHH**SSGLVPRGSHMGSLSEYQKLAQKKHEEEEEESYSHAMQLAMGVVLPMATQAAIQLGVF
EIIAKAGELSAPEIAAQLQAQNVKAPMMLDRMLRLLVSHRVLECSVSGGERLYALNPVSKYFVSNKDG
ASLGHFMALPLDKVFMESWLGLKDAVMEGGIPFNRVHGMHIFEYASGNPRFNETYHEAMFNHSTIAME
RILEHYEGFQNVERLVDVGGGFGVTLSMITSKYPQIKAVNFDLPHVVQDAPSYAGVEHVGGNMFESVP
EGDAILMKWILHCWDDDHCLRILKNCYKAVPGNGKVIVMNSIVPEIPEVSSAARETSLLDVLLMTRDG
GGRERTKKEYTELAIAAGFKGINFASCVCNLYIMEFFK

>*Cs*CaOMT (41.2 kDa)

MGSS**HHHHHH**SSGLVPRGSHMDSIVDGERDQSFAYASQLVMGTVLPMAIQAVYELGIFEILDKVGPGA
KLCASDIAAQLLTKNKDAPMMLDRILRLLASYSVVECSLDASGARRLYSLNSVSKYYVPNKDGVLLGP
LLQMNQDKVLLESWSQLKDAILEGGIPFNRAHGVHVFEYAGLDPKFNKHFNTAMYNYTSLVMSNILES
YKGFDNIKQLVDVGGSLGITLQAITTKYPYIKGINFDQPHVIDHAPSHPRIEHVGGDMFQSVPKGDAI
IMKSVLHDWNDEHCLKLLKNCYKSIPEDGKVIVVESMLPEVPNTSIESKSNSHFDVLMMIQSPGGKER
TRHEFMTLATGAGFGGISCELAIGNLWVMEFYK

>*Ms*CaOMT (42.1 kDa)

MGSS**HHHHHH**SSGLVPRGSHMGSTGETQITPTHISDEEANLFAMQLASASVLPMILKSALELDLLEII
AKAGPGAQISPIEIASQLPTTNPDAPVMLDRMLRLLACYIILTCSVRTQQDGKVQRLYGLATVAKYLV
KNEDGVSISALNLMNQDKVLMESWYHLKDAVLDGGIPFNKAYGMTAFEYHGTDPRFNKVFNKGMSDHS
TITMKKILETYTGFEGLKSLVDVGGGTGAVINTIVSKYPTIKGINFDLPHVIEDAPSYPGVEHVGGDM
FVSIPKADAVFMKWICHDWSDEHCLKFLKNCYEALPDNGKVIVAECILPVAPDSSLATKGVVHIDVIM
LAHNPGGKERTQKEFEDLAKGAGFQGFKVHCNAFNTYIMEFLKKV

> KDO86634.1 (*Citrus sinensis*) (45.7 kDa)

MGSS**HHHHHH**SSGLVPRGSHMGSLSEYQKLAQKKHEEEEEEEESYSHAMQLAMGVVLPMATQAAIQLG
VFEIIAKAGELSAPEIAAQLQAQNVKAPMMLDRMLRLLVSHRVLECSVSGGERLYALNPVSKYFVSNK
DGASLGHFMALPLDKVFMESWYIIILSFFFFPLSGQIYIVVNLSNFKNACRLGLKDAVMEGGIPFNRV
HGMHIFEYASGNPRFNETYHEAMFNHSTIAMERILEHYEGFQNVERLVDVGGGFGVTLSMITSKYPQI
KAVNFDLPHVVQDAPSYAGVEHVGGNMFESVPEGDAILMKWILHCWDDDHCLRILKNCYKAVPGNGKV
IVMNSIVPEIPEVSSAARETSLLDVLLMTRDGGGRERTKKEYTELAIAAGFKGINFASCVCNLYIMEF
FK

### Supplementary Figures

#### SDS-PAGE analysis

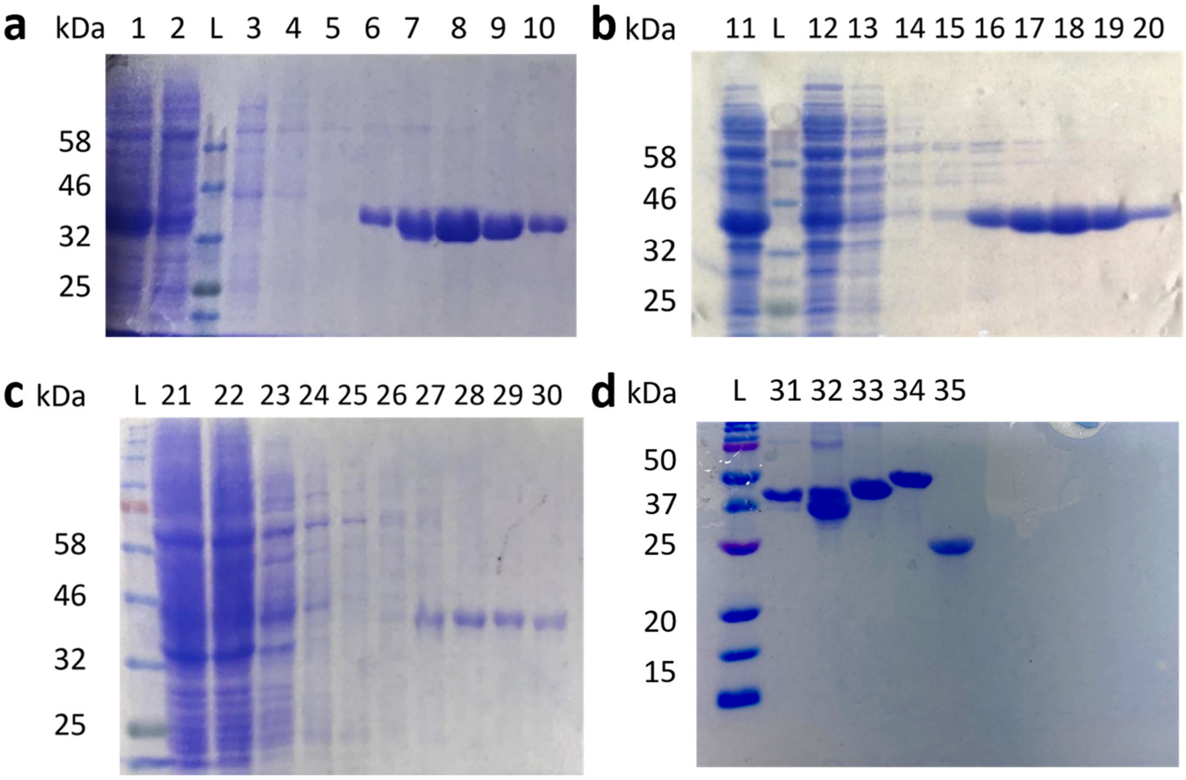

**Figure S 1** – Visualisation of the enzymes used in this study. L=Ladder. **a**: SDS gel of *Cs*ANMT (41.9 kDa) fractions during purification. 1: crude lysate; 2: flow through; 3: 5 mM imidazole; 4: 10 mM imidazole; 5: 20 mM imidazole; 6: 50 mM imidazole; 7: 100 mM imidazole; 8: 150 mM imidazole; 9: 200 mM imidazole; 10: 300 mM imidazole. **b**: SDS gel of *Cs*CaOMT (41.2 kDa) fractions during purification. 11: crude lysate; 12: flow through; 13: 5 mM imidazole; 14: 10 mM imidazole; 15: 20 mM imidazole; 16: 50 mM imidazole; 17: 100 mM imidazole; 18: 150 mM imidazole; 19: 200 mM imidazole; 20: 300 mM imidazole. **c**: SDS gel of *Ms*CaOMT (42.1 kDa) fractions during purification. 21: crude lysate; 22: flow through; 23: 5 mM imidazole; 24: 10 mM imidazole; 25: 20 mM imidazole; 26: 50 mM imidazole; 27: 100 mM imidazole; 28: 150 mM imidazole; 29: 200 mM imidazole; 30: 300 mM imidazole. **d**: 31: *Rg*ANMT (42.2 kDa); 32: *Pp*CaOMT (43.7 kDa); 33: *Ec*MAT (44.1 kDa); *Tk*MAT (46.7 kDa); *Ec*MTAN (26.5 kDa).

#### Amino acid alignment

```
CLUSTAL O(1.2.4) multiple sequence alignment

MsCaOMT             --MGSTGETQITP------------THISDEEA----NLFAMQL-------ASASVLPMI    35
RgANMT              --MGSLSES------HTQYKHG-VEVEEDEEES----YSRAMQL-------SMAIVLPMA    40
CsANMT              --MGSLSEY------QKLAQKK-HEEE--EEES----YSHAMQL-------AMGVVLPMA    38
KDO86634.1(Cs)      --MGSLSEY------QKLAQKK-HEEEEEEEES----YSHAMQL-------AMGVVLPMA    40
PpCaOMT             --MASSLERKSHPKINHAEPED-EITKEEEDES----FCYAMQL-------VGSSVLSMS    46
CsCaOMT             --------------------MD-SIVDGERDQS----FAYASQL-------VMGTVLPMA    28
RnCOMT              --MGDTKEQRILRYVQQNAKPGDPQSVLEAIDTYCTQKEWAMNVGDAKGQIMDAVIREYS    58
MxSafC              MIHHVELTQSVLQYIRDSSVRD-NDILRDLREETSKLPLRTMQIPPEQGQLLSLLVRLIG    59
                                                   :        : ::           :

MsCaOMT             LKSALELDLLEIIAKAGPGAQISPIEIASQLPTTNPDAPVMLDRMLRLLACYIILTCSVR    95
RgANMT              TQSAIQLGVFEIIAK-APGGRLSASEIATILQAQNPKAPVMLDRMLRLLVSHRVLDCSVS    99
CsANMT              TQAAIQLGVFEIIAK-A--GELSAPEIAAQLQAQNVKAPMMLDRMLRLLVSHRVLECSVS    95
KDO86634.1(Cs)      TQAAIQLGVFEIIAK-A--GELSAPEIAAQLQAQNVKAPMMLDRMLRLLVSHRVLECSVS    97
PpCaOMT             LQSAIKLGIFDIIARKGPGAKLSSSEIATKIGTENPEAPVMVDRILRLLTSHSVLNCSAV    106
CsCaOMT             IQAVYELGIFEILDKVGPGAKLCASDIAAQLLTKNKDAPMMLDRILRLLASYSVVECSLD    88
RnCOMT              PSLVLELGAYCGYSA---------VRMA-----------------RLLQPGARLLTMEMN    92
MxSafC              ARKTLEVGVFTGYST---------LCAA-----------------LALPADGRVIACDLS    93
                       . ::.                   *                   *     ::  .

MsCaOMT             TQQDG-KVQRLYGLATVA-KYLVKNEDG-----VSISALNLMNQDKVLMESWY-------    141
RgANMT              G----PAGERLYGLTSVS-KYFVPDQDG-----ASLGNFMALPLDKVFMESWM-------    142
CsANMT              G------GERLYALNPVS-KYFVSNKDG-----ASLGHFMALPLDKVFMESWL-------    136
KDO86634.1(Cs)      G------GERLYALNPVS-KYFVSNKDG-----ASLGHFMALPLDKVFMESWYIIILSFF    145
PpCaOMT             AANGGSDFQRVYSLGPVS-KYFVNDEEG-----GSLGPLLTLIQDRVFLESWS-------    153
CsCaOMT             A----SGARRLYSLNSVS-KYYVPNKDG-----VLLGPLLQMNQDKVLLESWS-------    131
RnCOMT              PDYAA-ITQQMLNFAGLQDKVTILNGASQDLIPQLKKKYDVDTLDMVFLDHWKDRYLPDT    151
MxSafC              EEWVS-IARRYWQRAGVADRIEVRLGDAHHSLEALVGSEHRGTFDLAFIDADKESYDF--    150
                            .:      :  :  :    .                * .:::

MsCaOMT             -----------------------HLKDAVLDGGIPFNKAYGMTAFEYHGTDPRFNKVFNK    178
RgANMT              -----------------------GVKGAVMEGGIPFNRVHGMHIFEYASSNSKFSDTYHR    179
CsANMT              -----------------------GLKDAVMEGGIPFNRVHGMHIFEYASGNPRFNETYHE    173
KDO86634.1(Cs)      FFPLSGQIYIVVNLSNFKNACRLGLKDAVMEGGIPFNRVHGMHIFEYASGNPRFNETYHE    205
PpCaOMT             -----------------------QLKDAVVEGGIPFNRVHGMHAFEYPGLDPRFNQVFNT    190
CsCaOMT             -----------------------QLKDAILEGGIPFNRAHGVHVFEYAGLDPKFNKHFNT    168
RnCOMT              ---------------LLLEKCGLLRKG---------------------------------    163
MxSafC              ---------------YYEHALRLVRPG---------------------------------    162
                                              .

MsCaOMT             GMSDHSTITMKKILETYTGFE--GLKSLVDVGGGTGAVINTIVSKYPTIKGINFDLPHVI    236
RgANMT              AMFNHSTIALKRILEHYKGFE--NVTKLVDVGGGLGVTLSMIASKYPHIQAINFDLPHVV    237
CsANMT              AMFNHSTIAMERILEHYEGFQ--NVERLVDVGGGFGVTLSMITSKYPQIKAVNFDLPHVV    231
KDO86634.1(Cs)      AMFNHSTIAMERILEHYEGFQ--NVERLVDVGGGFGVTLSMITSKYPQIKAVNFDLPHVV    263
PpCaOMT             AMFNHTTIVIKKLLHIYKGLEDKNLTQLVDVGGGLGVTLNLITSRYQHIKGINFDLPHVV    250
CsCaOMT             AMYNYTSLVMSNILESYKGFD--NIKQLVDVGGSLGITLQAITTKYPYIKGINFDQPHVI    226
RnCOMT              ------------------------------------------------------------    163
MxSafC              ------------------------------------------------------------    162

MsCaOMT             EDAPSYPGVEHVGGDMFVSIPKADAVFMKWICHDWSDEHCLKFLKNCYEALPDNGKVIVA    296
RgANMT              QDAASYPGVEHVGGNMFESVPEGDAILMKWILHCWDDEQCLRILKNCYKATPENGKVIVM    297
CsANMT              QDAPSYAGVEHVGGNMFESVPEGDAILMKWILHCWDDDHCLRILKNCYKAVPGNGKVIVM    291
KDO86634.1(Cs)      QDAPSYAGVEHVGGNMFESVPEGDAILMKWILHCWDDDHCLRILKNCYKAVPGNGKVIVM    323
PpCaOMT             NHAPSYPGVEHVGGDMFASVPSGDAIFMKWILHDWSDEHCLKLLKNCYKAIPDNGKVIVV    310
CsCaOMT             DHAPSHPRIEHVGGDMFQSVPKGDAIIMKSVLHDWNDEHCLKLLKNCYKSIPEDGKVIVV    286
RnCOMT              ------------------------------------------------------TVLLAD    169
MxSafC              ------------------------------------------------------GLIILD    168
                                                                        ::
```

```
MsCaOMT             ECILPVAPDSSLATKGVVHIDVIMLAHNPGGKERTQKEFEDLAKGAGFQGFKVHCNAFNT    356
RgANMT              NSVVPETPEVSSSARETSLLDVLLMTRDGGGRERTQKEFTELAIGAGFKGINFACCVCNL    357
CsANMT              NSIVPEIPEVSSAARETSLLDVLLMTRDGGGRERTKKEYTELAIAAGFKGINFASCVCNL    351
KDO86634.1(Cs)      NSIVPEIPEVSSAARETSLLDVLLMTRDGGGRERTKKEYTELAIAAGFKGINFASCVCNL    383
PpCaOMT             EALLPAMPETSTATKTTSQLDVLMMTQNPGGKERSEQEFMALATGAGFSGIRYECFVCNF    370
CsCaOMT             ESMLPEVPNTSIESKSNSHFDVLMMIQSPGGKERTRHEFMTLATGAGFGGISCELAIGNL    346
RnCOMT              NVIVPGTPDFLAYVRGSSS--F------ECTHYSSYLEYMKVVDGLEKAIYQGPSSPDKS    221
MxSafC              NTLWSGKVADPSVVGDPETDSLRRINAKLLTDERVDLSMLPIADGLTLARKR--------    220
                    : :                  .               .   :. .

MsCaOMT             YIMEFLKKV  365
RgANMT              HIMEFFK--  364
CsANMT              YIMEFFK--  358
KDO86634.1(Cs)      YIMEFFK--  390
PpCaOMT             WVMEFFK--  377
CsCaOMT             WVMEFYK--  353
RnCOMT              ---------  221
MxSafC              ---------  220
```

**Figure S 2 –** Amino acid alignment of all used MTs (*Rg*ANMT; *Cs*ANMT; KDO86634.1 (*Cs*); *Cs*CaOMT; *Pp*CaOMT; *Ms*CaOMT) and additionally *Rn*COMT and *Mx*SafC as catechol *O*-MTs (class I enzymes according to *Joshi et al.*) for comparison.[6] The alignment was created by Clustal Omega.[7,8]

#### Enzyme screening

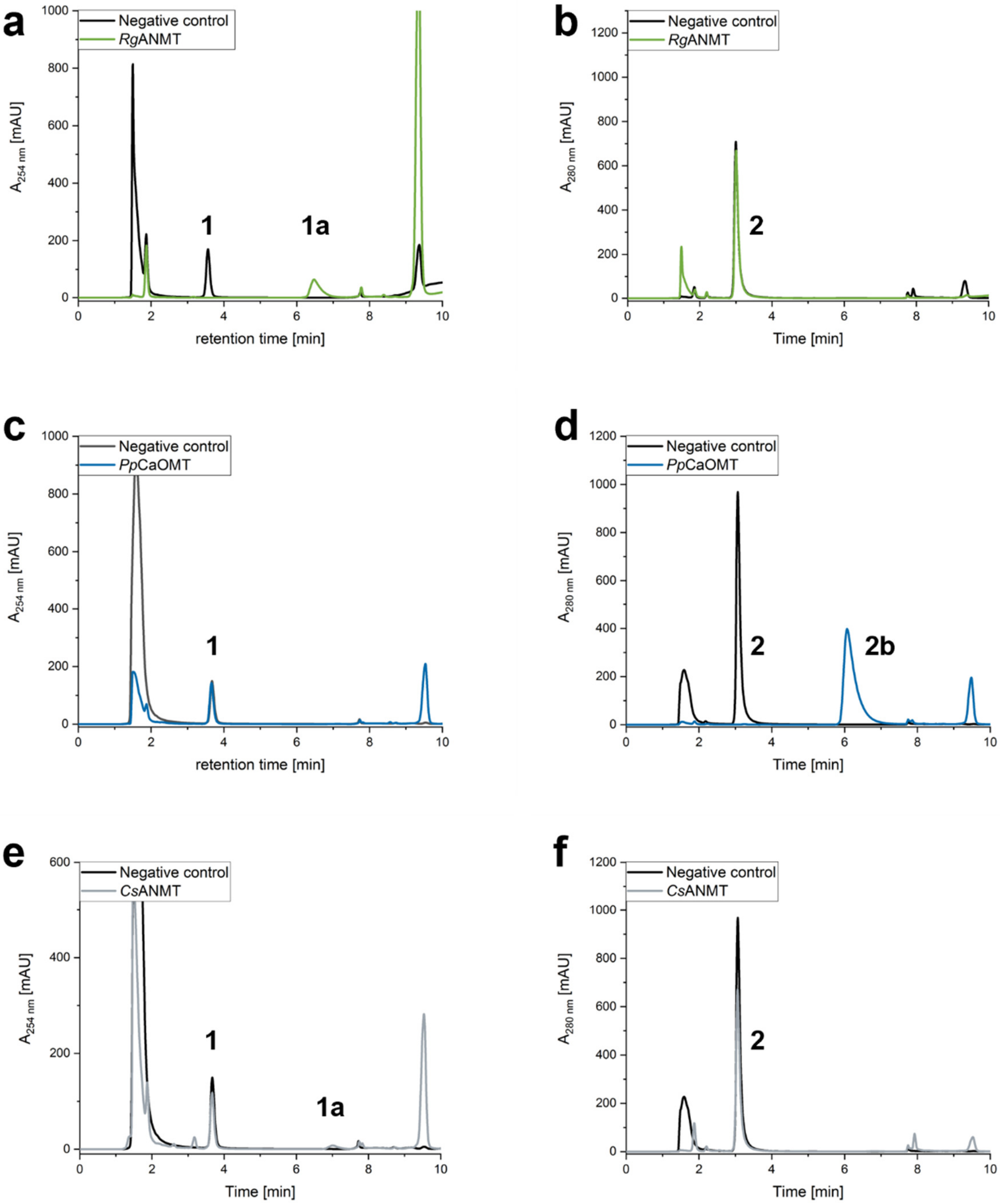

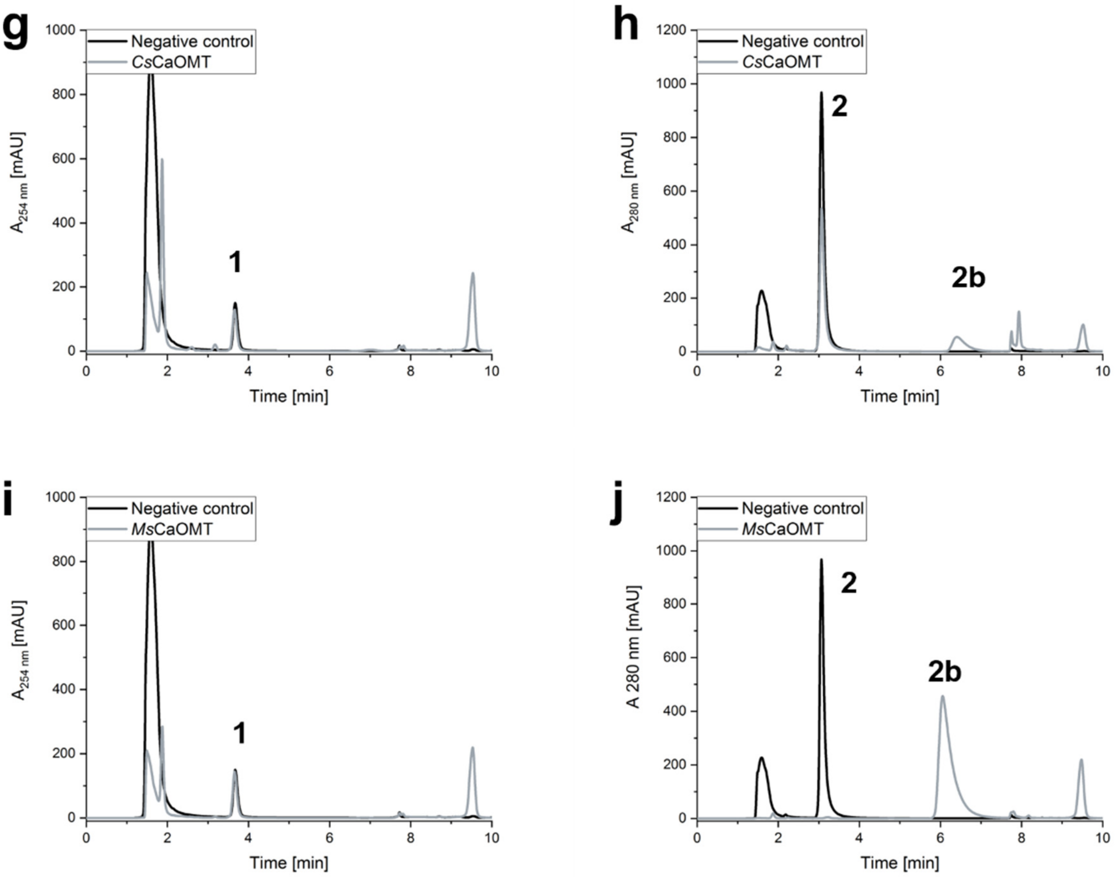

**Figure S 3** – Enzyme screening analysed by HPLC Method A using the natural substrates **1** for the confirmation of *N*-methylation and **2** for *O*-methylation. The negative controls did not contain an MT but the MAT and MTAN enzymes (black). **a – b**: Reactions catalysed by *Rg*ANMT. **1** was fully converted to **1a** while **2** was not accepted as substrate. **c – d**: Reactions catalysed by *Pp*CaOMT. **1** was not accepted while **2** was fully converted to **2b. e – f**: *Cs*ANMT accepted **1** as substrate and partly formed **1a**. **2** was not accepted as substrate. **g – h**: *Cs*CaOMT only accepted **2** but not **1** as substrate and parly produced **2b**. **i – j**: *Ms*CaOMT only accepted **2** as substrate leading to full conversion to product **2b.** This was already shown before.[9]

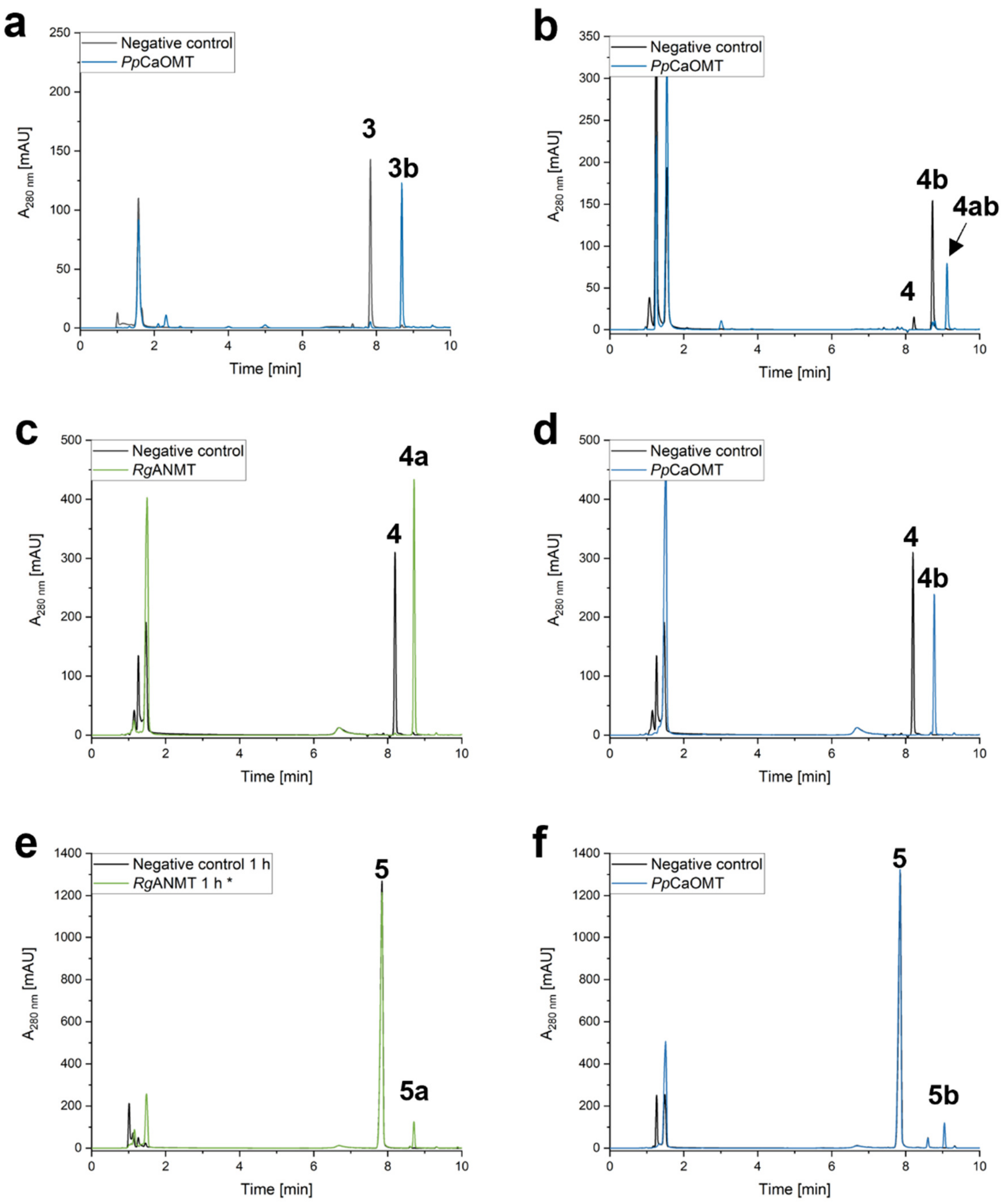
a
Negative control
PpCaOMT
3
3b
A280 nm [mAU]
Time [min]
b
Negative control
PpCaOMT
4
4b
4ab
c
Negative control
RgANMT
4
4a
d
Negative control
PpCaOMT
4
4b
e
Negative control 1 h
RgANMT 1 h *
5
5a
f
Negative control
PpCaOMT
5
5b

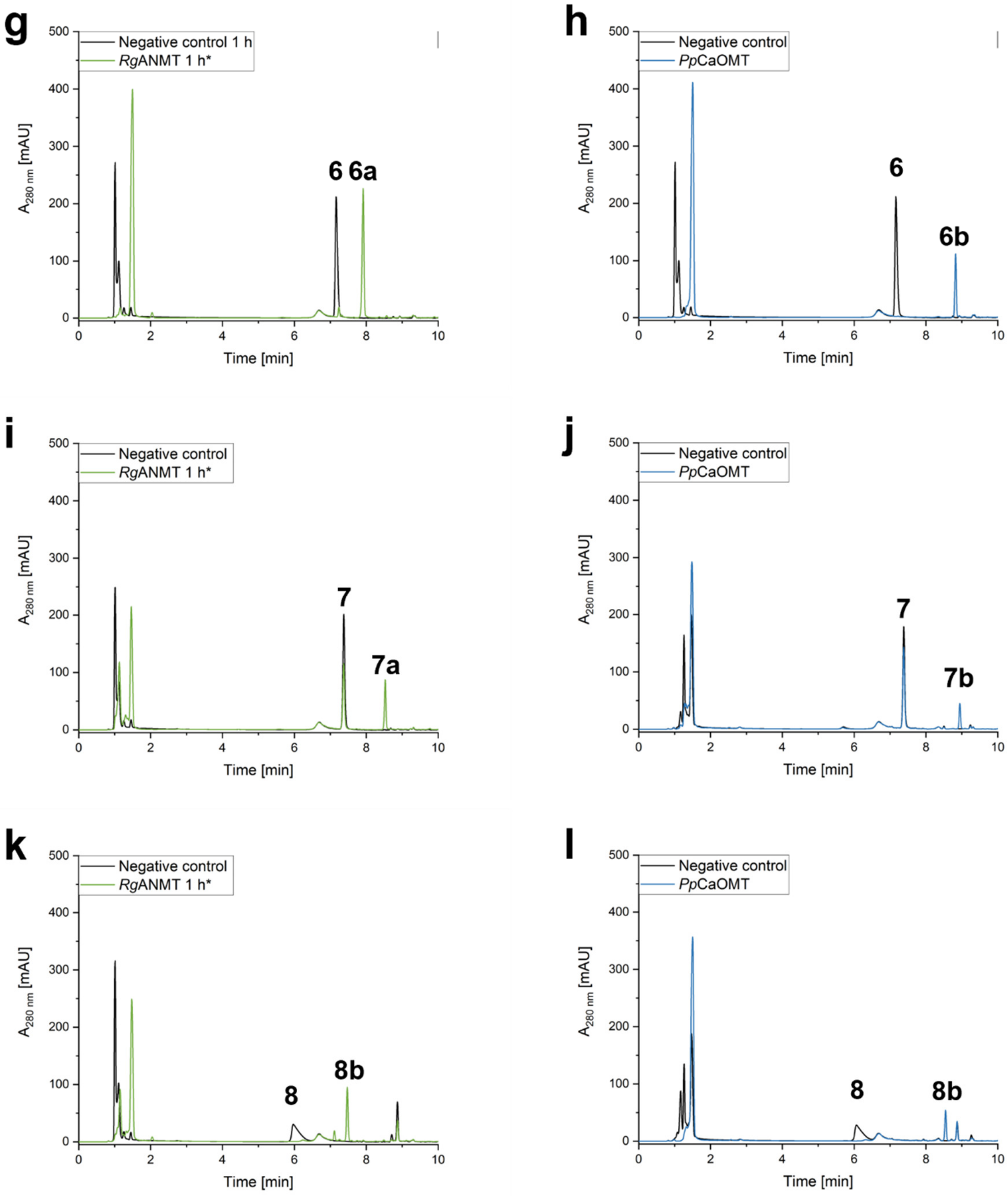

**Figure S 4** – HPLC chromatograms (Method B) of the substrate screening using *Rg*ANMT and *Pp*CaOMT. Samples were taken after 20 h. The *N*-methylated products for substrate **5** – **8** were not stable under assay conditions and the product peaks were decreased after 20 h compared to the 1 h samples. The formation of **5a - 8a** are shown after 1 h . **a**: Conversion of **3** to **3b** catalysed by *Pp*CaOMT. **b**: Product **4a** was used as substrate for another methylation step catalysed by *Pp*CaOMT to form product **4ab**. **4a** was synthesised in vivo. Small amounts of the origin substrate **4** were left in the stock solution leading to the formation of **4b** in small amounts as well. **c**: Conversion from **4** to **4a** catalysed by *Rg*ANMT. **d**: Conversion of **4** to **4b** catalysed by *Pp*CaOMT. **e**: Formation of **5a** catalysed by *Rg*ANMT after 1 h. **f**: Formation of **5b** catalysed by *Pp*CaOMT. **g**: Formation of **6a** catalysed by *Rg*ANMT after 1 h. **h**: Formation of **6b** catalysed by *Pp*CaOMT. **i**: Formation of **7a** catalysed by *Rg*ANMT after 1 h. **j**: Formation of **7b** catalysed by *Pp*CaOMT. **k**: Formation of **8a** catalysed by *Rg*ANMT after 1 h. **l**: Formation of **8b** catalysed by *Pp*CaOMT.

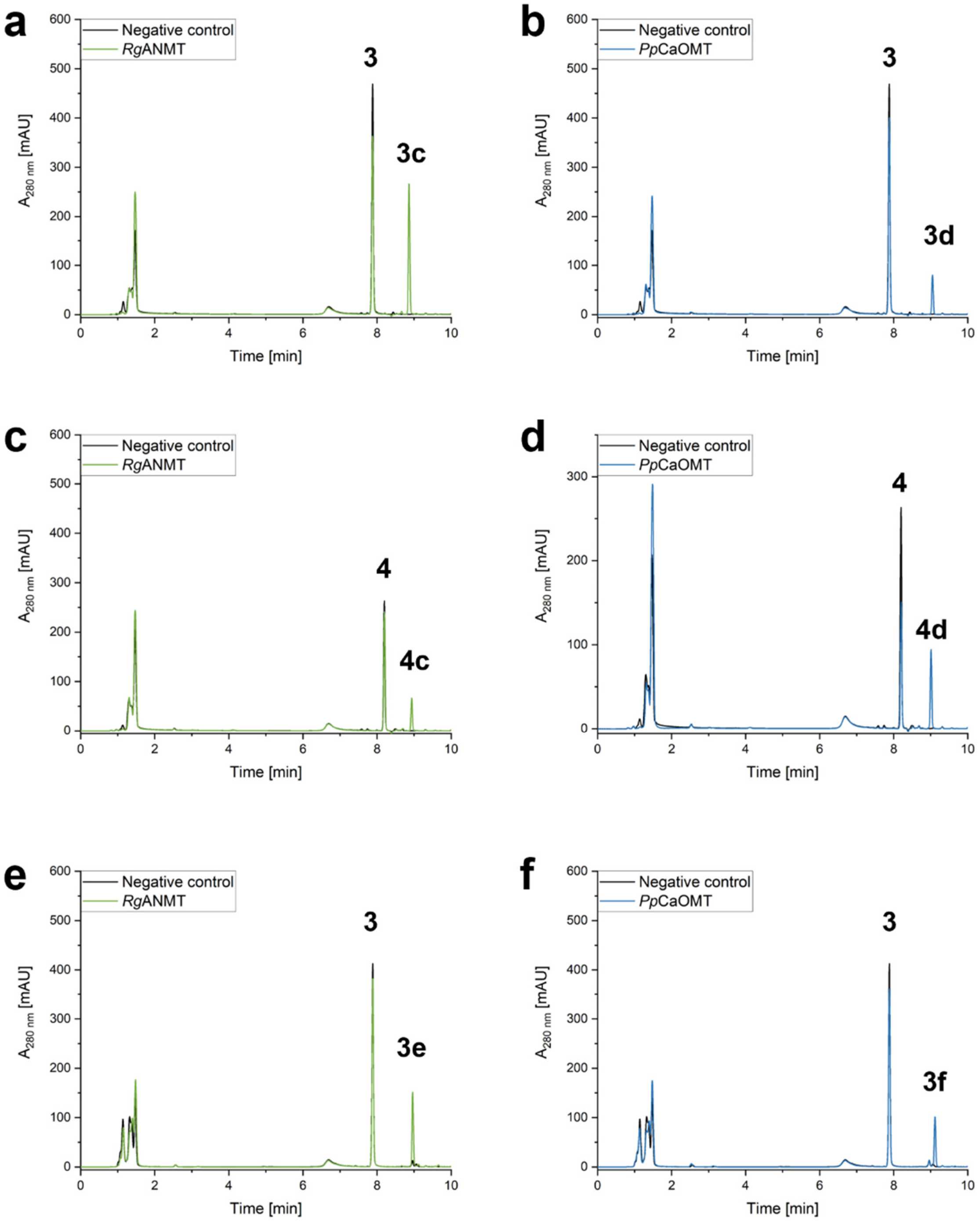

a
b
c
d
e
f
Negative control
RgANMT
PpCaOMT
3
3c
3d
4
4c
4d
3e
3f
A280 nm [mAU]
Time [min]

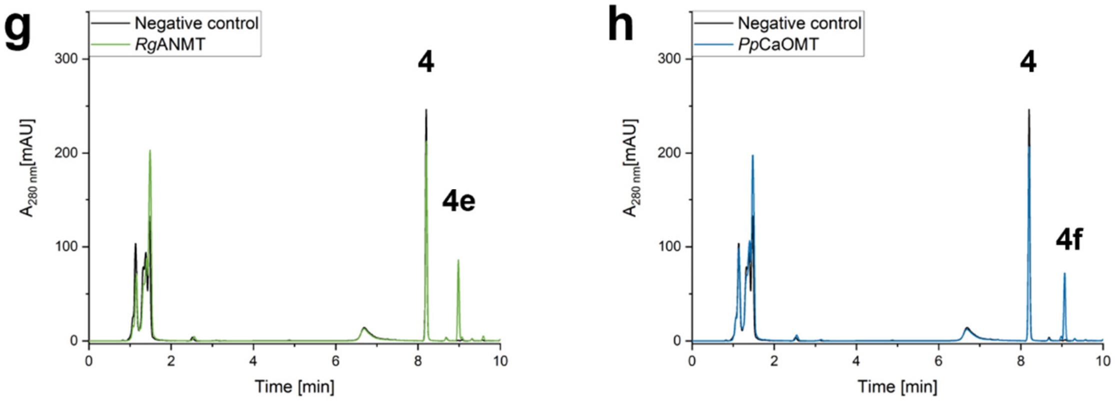

**Figure S 5** – HPLC chromatograms (Method B) of the ethyl and allyl transfer assays using *Rg*ANMT and *Pp*CaOMT as biocatalysts. The samples were taken after 20 h. **a**: Formation of **3c** catalysed by *Rg*ANMT. **b**: Formation of **3d** catalysed by *Pp*CaOMT. **c**: Formation of **4c** catalysed by *Rg*ANMT. **d**: Formation of **4d** catalysed by *Pp*CaOMT. **e**: Formation of **3e** catalysed by *Rg*ANMT. **f**: Formation of **3f** catalysed by *Pp*CaOMT. **g**: Formation of **4e** catalysed by *Rg*ANMT. **h**: Formation of **4f** catalysed by *Pp*CaOMT.

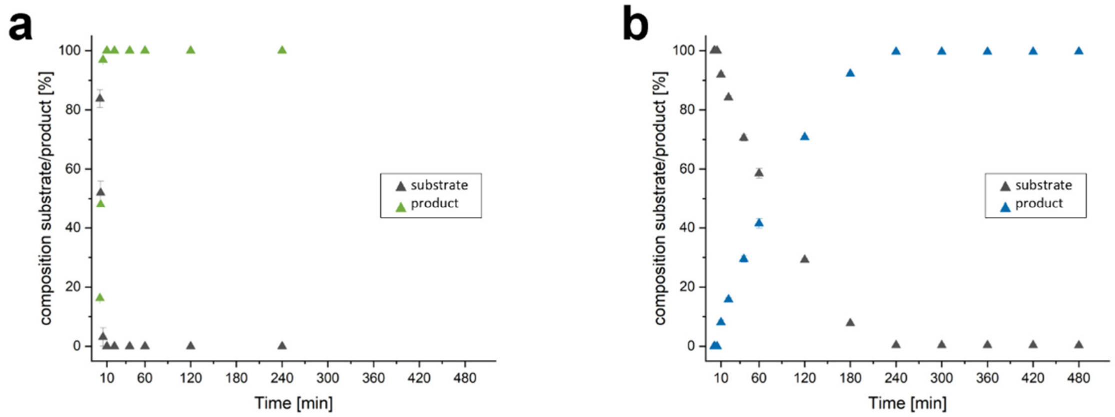

**Figure S 6** – Time course experiments with *Rg*ANMT and *Pp*CaOMT using substrate **4**. **a**: Reaction catalysed by *Rg*ANMT. After 5 min the reaction is completed and no substrate is left in the assay. **b**: The reaction catalysed by *Pp*CaOMT with substrate **4** is completed after 240 min.

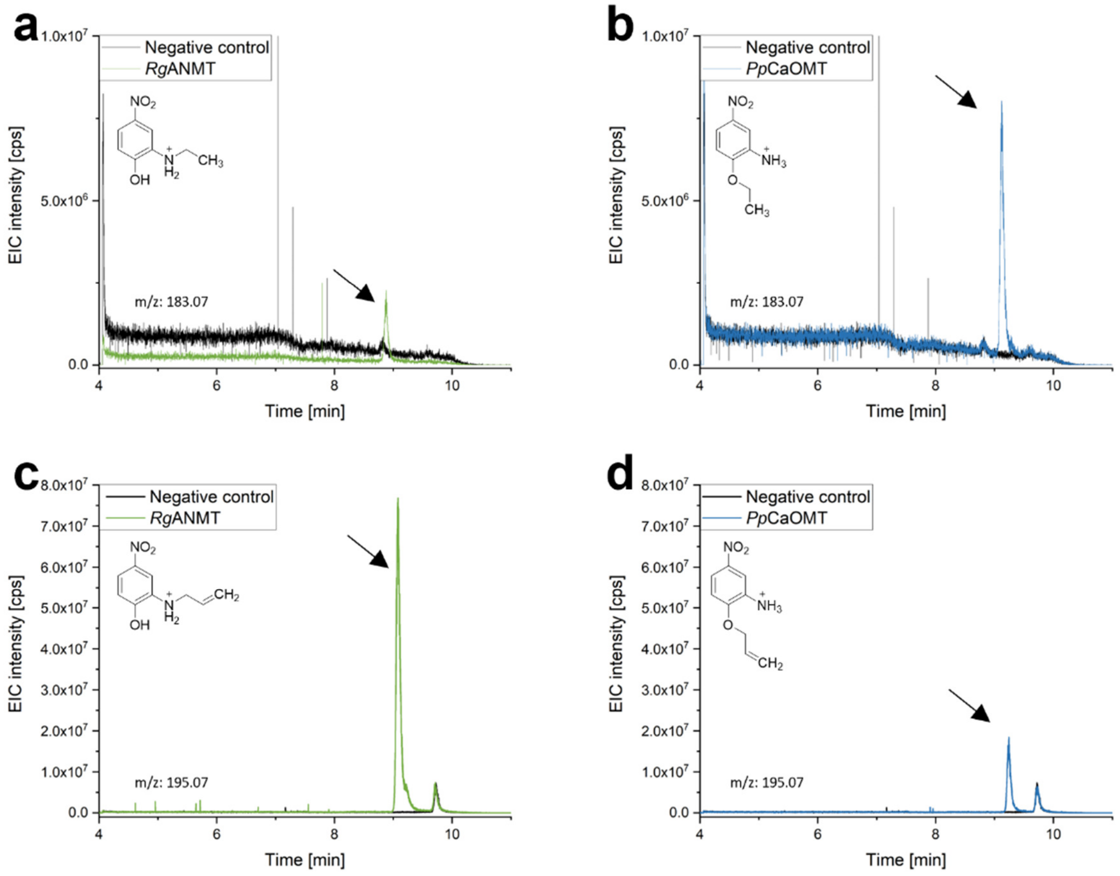

**Figure S 7** – Extracted ion chromatograms for ethylation and allylation reactions catalysed by *Rg*ANMT and *Pp*CaOMT. The product was found in the positive mode. **a**: Formation of **3c** catalysed by *Rg*ANMT. **b**: Formation of **3d** catalysed by *Pp*CaOMT. **c**: Formation of **3e** catalysed by *Rg*ANMT. **c**: Formation of **3f** catalysed by *Pp*CaOMT.

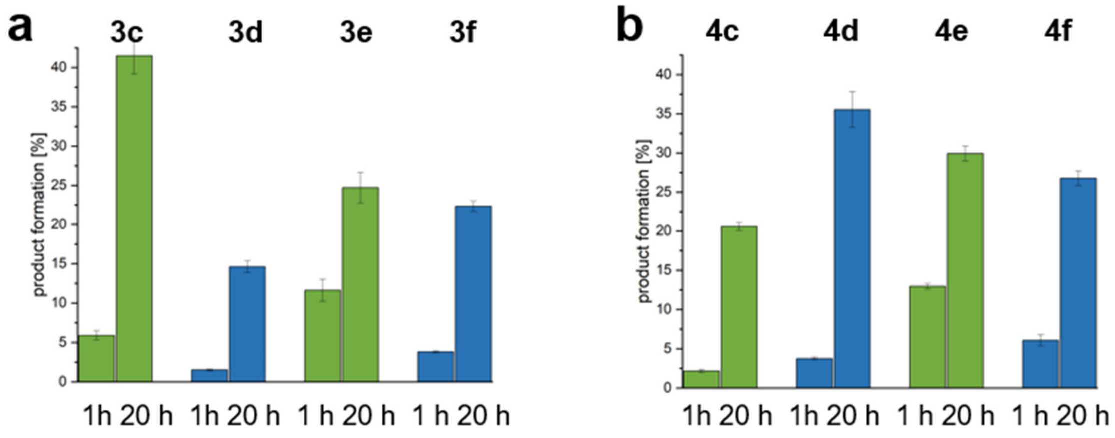

**Figure S 8** – Conversion numbers for ethylation and allylation experiments catalysed by *Rg*ANMT (green) and *Pp*CaOMT (blue) after 1 and 20 h. **a**: Conversion of ethylated and allylated products **3c**-**f** using substrate **3**. **b**: Conversion of ethylated and allylated products **4c**-**f** using substrate **4**.

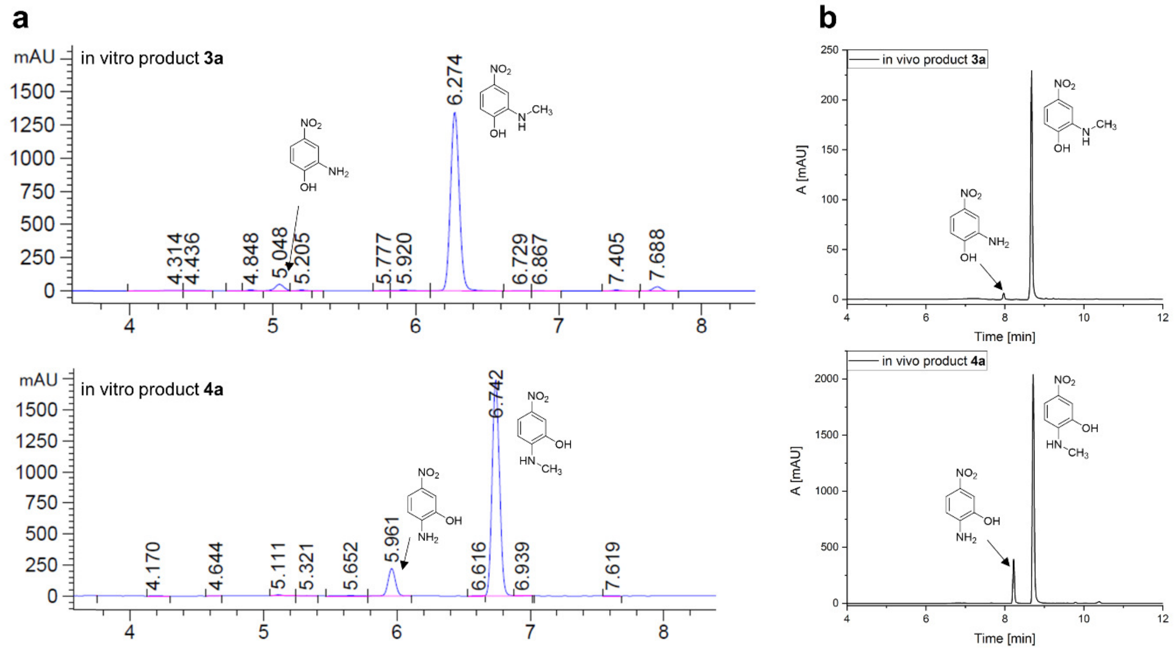

**Figure S 9** – HPLC chromatograms of purified *N*-methylated products **3a** and **4a**. **a**: Products from upscale in vitro experiments catalysed by *Rg*ANMT. **b**: Products from upscale in vivo experiments catalysed by *Rg*ANMT.

### NMR analysis

#### Confirmation of chemoselective reactions

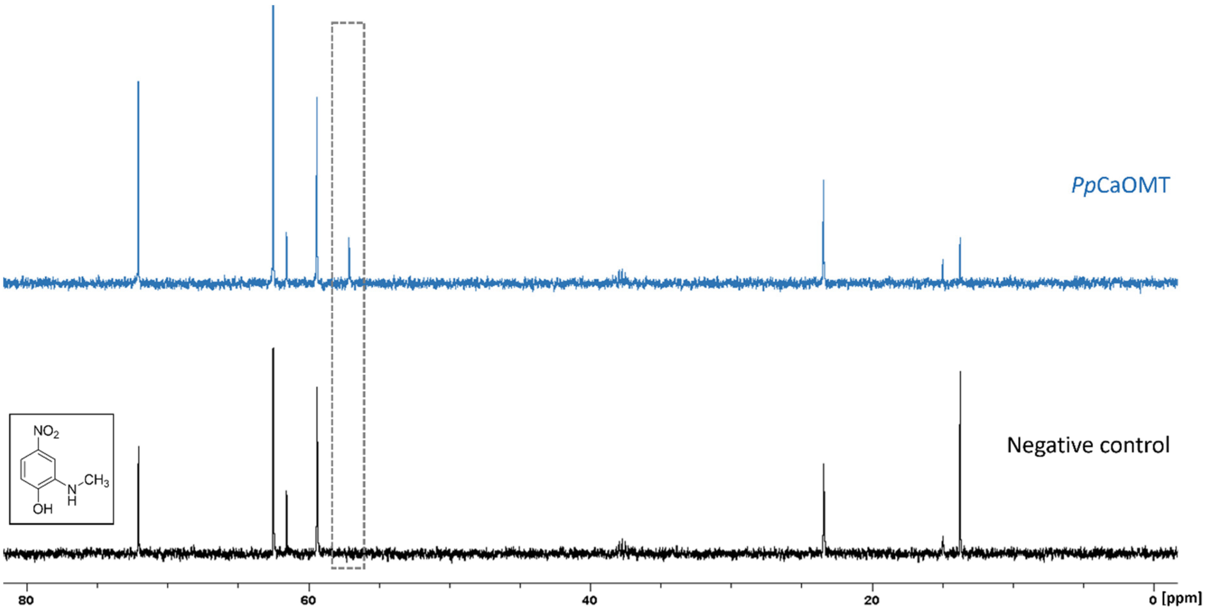

**Figure S 10** – Double methylation using product **3a** as substrate. For the exeriment, $^{13}$C-labelled methionine was used. The concentration of **3a** was too low to visualize the carbon of the methyl group in *N*-position. The signal in the *Pp*CaOMT reaction (blue) at 58 ppm belongs to the $^{13}$C-labelled methyl group adjacent to the hydroxyl group. $^{13}$C-NMR (100.6 MHz, 5% $D_2O$); all other signals are assigned in **Table S 5** in the NMR chapter.

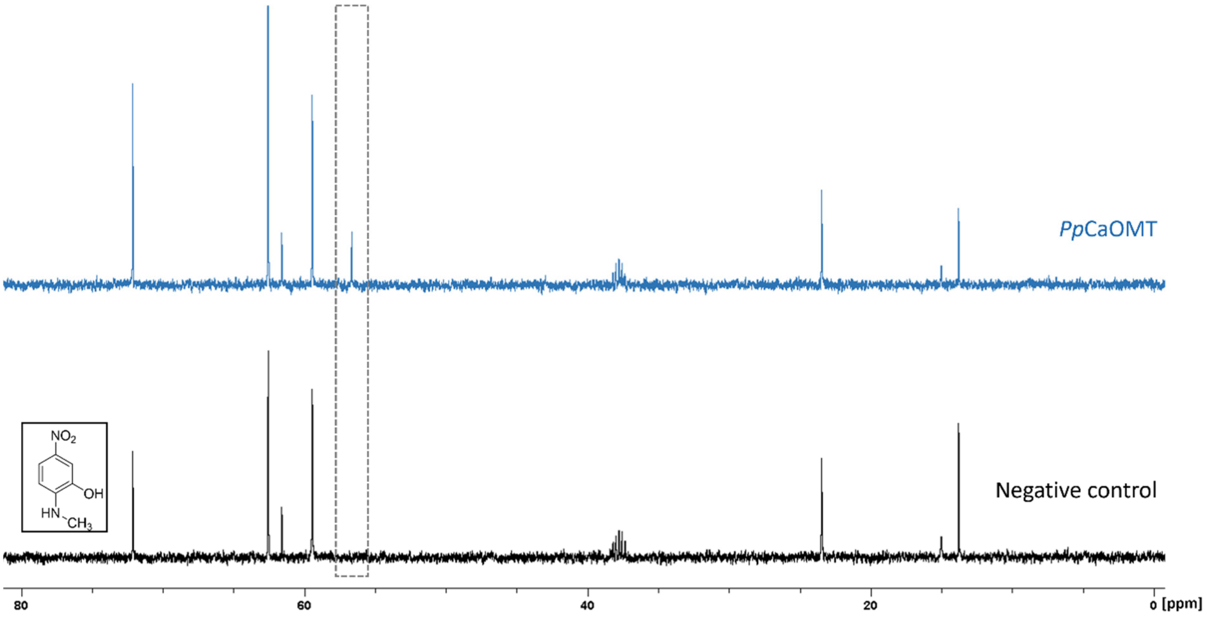

**Figure S 11** – Double methylation using product **4a** as substrate. For the exeriment, $^{13}$C-labelled methionine was used. The concentration of **4a** was too low to visualize the carbon of the methyl group in *N*-position. The signal in the *Pp*CaOMT reaction (blue) at 58 ppm belongs to the $^{13}$C-labelled methyl group adjacent to the hydroxyl group. $^{13}$C-NMR (100.6 MHz, 5% $D_2O$); all other signals are assigned in **Table S 5** in the NMR chapter.

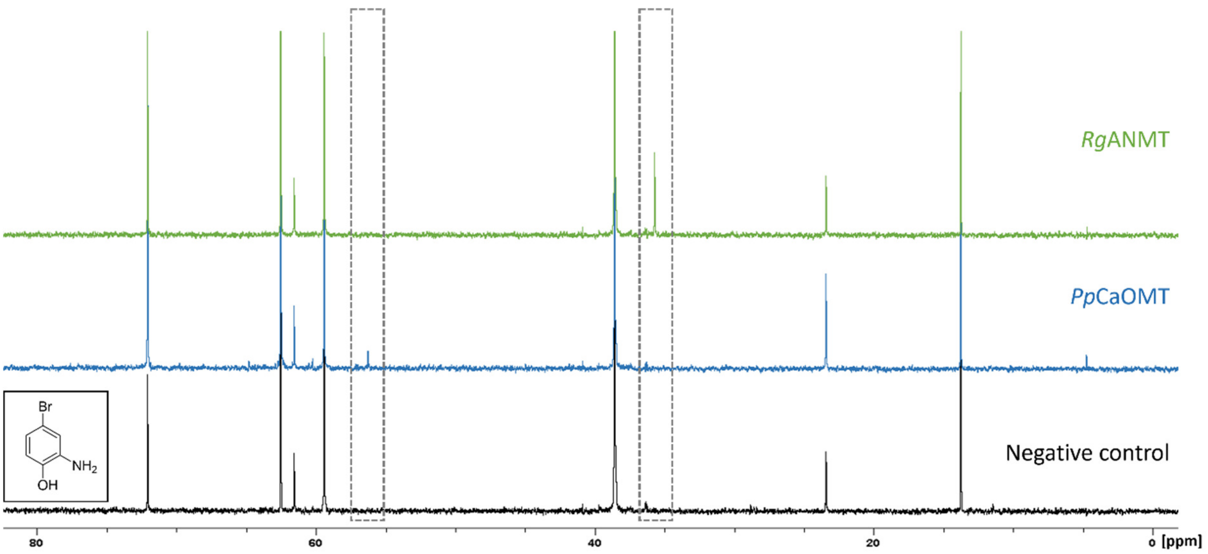

**Figure S 12** – Methylation reaction catalysed by *Rg*ANMT (green) and *Pp*CaOMT (blue) using substrate **5**. The $^{13}C$-labelled methyl group was transferred onto the amino group (signal at 32 ppm) in the *N*-MT reaction and onto the hydroxyl group (signal at 56 ppm) in the *O*-MT reaction. $^{13}C$-NMR (100.6 MHz, 5% $D_2O$); all other signals are assigned in **Table S 5** in the NMR chapter.

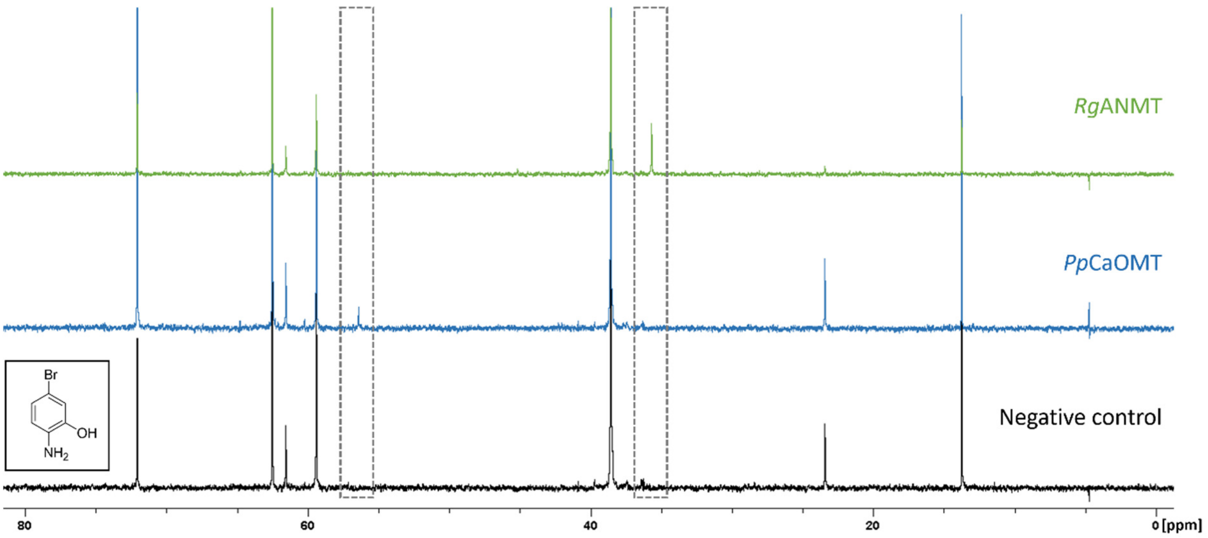

**Figure S 13** – Methylation reaction catalysed by *Rg*ANMT (green) and *Pp*CaOMT (blue) using substrate **6**. The $^{13}C$-labelled methyl group was transferred onto the amino group (signal at 31 ppm) in the *N*-MT reaction and onto the hydroxyl group (signal at 56 ppm) in the *O*-MT reaction. $^{13}C$-NMR (100.6 MHz, 5% $D_2O$); all other signals are assigned in **Table S 5** in the NMR chapter.

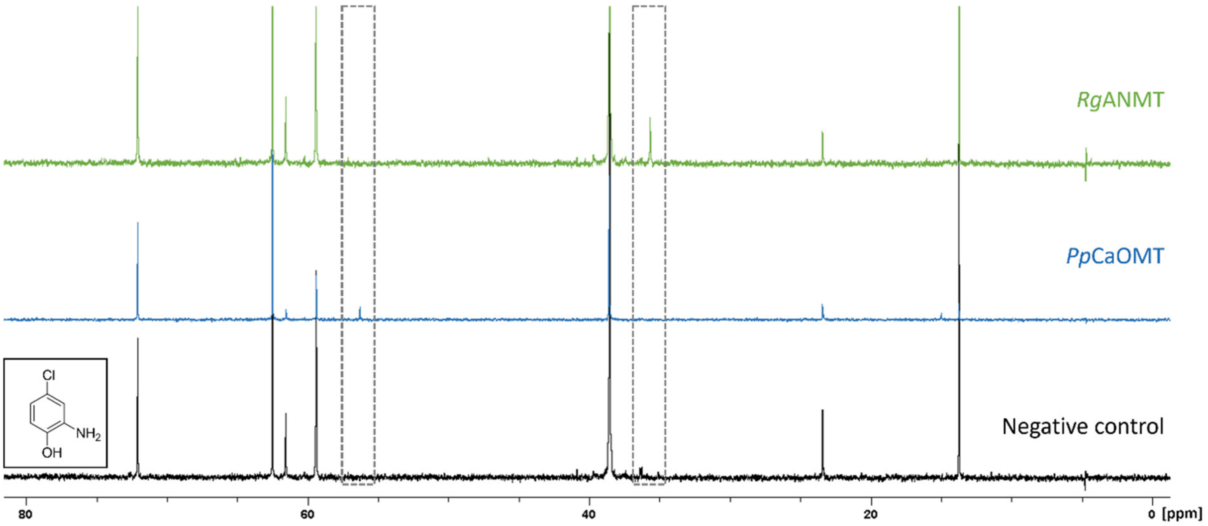

**Figure S 14** – Methylation reaction catalysed by *Rg*ANMT (green) and *Pp*CaOMT (blue) using substrate **7**. The $^{13}C$-labelled methyl group was transferred onto the amino group (signal at 31 ppm) in the *N*-MT reaction and onto the hydroxyl group (signal at 57 ppm) in the *O*-MT reaction. $^{13}C$-NMR (100.6 MHz, 5% $D_2O$); all other signals are assigned in **Table S 5** in the NMR chapter.

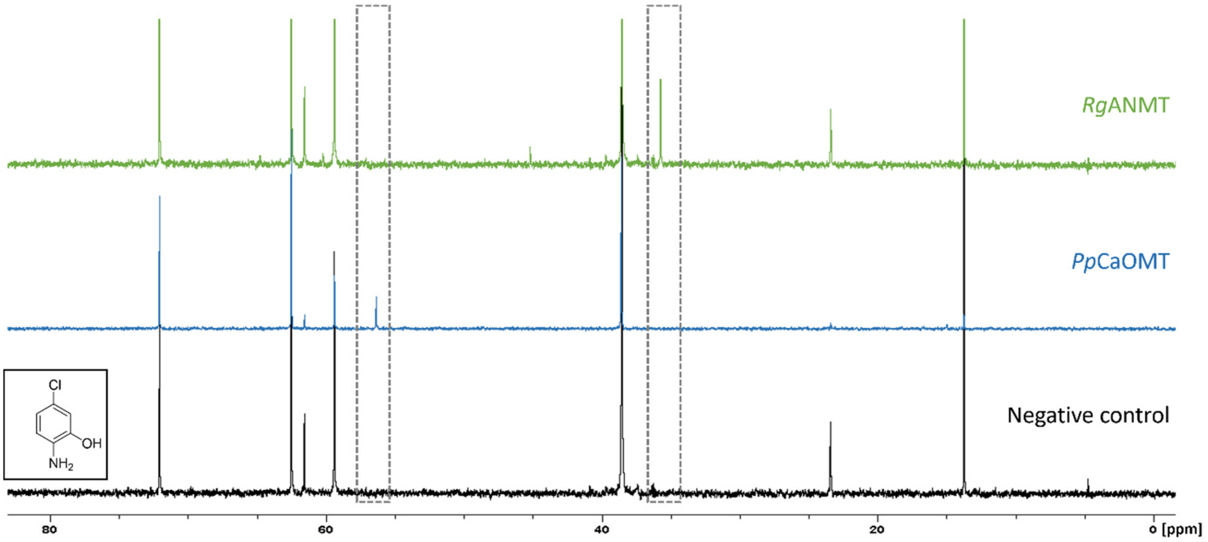

**Figure S 15** – Methylation reaction catalysed by *Rg*ANMT (green) and *Pp*CaOMT (blue) using substrate **8**. The $^{13}C$-labelled methyl group was transferred onto the amino group (signal at 31 ppm) in the *N*-MT reaction and onto the hydroxyl group (signal at 56 ppm) in the *O*-MT reaction. $^{13}C$-NMR (100.6 MHz, 5% $D_2O$); all other signals are assigned in **Table S 5** in the NMR chapter.

### Upscale reactions NMR

#### in vivo

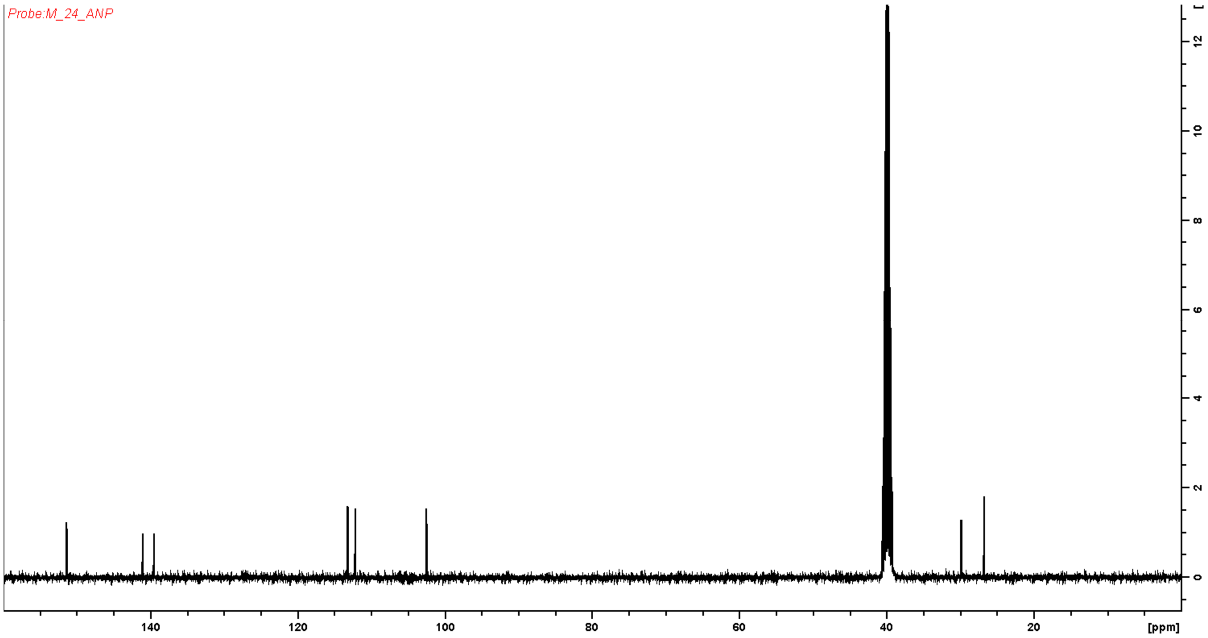

**Figure S 16** – $^{13}$C-NMR for product **3a** from in vivo upscale experiments catalysed *Rg*ANMT after purification. $^{13}$C-NMR (100.6 MHz, $D_6$MSO) δ(ppm): 151.5, 141.1, 139.8, 113.3, 112.4, 102.8, 29.9, 27.0. The signal at 27.0 ppm is assigned to cyclohexane which was left from the purification method with the puriflash. The signal at 29.9 was assigned to the carbon of the methyl group adjacent to the amino group.

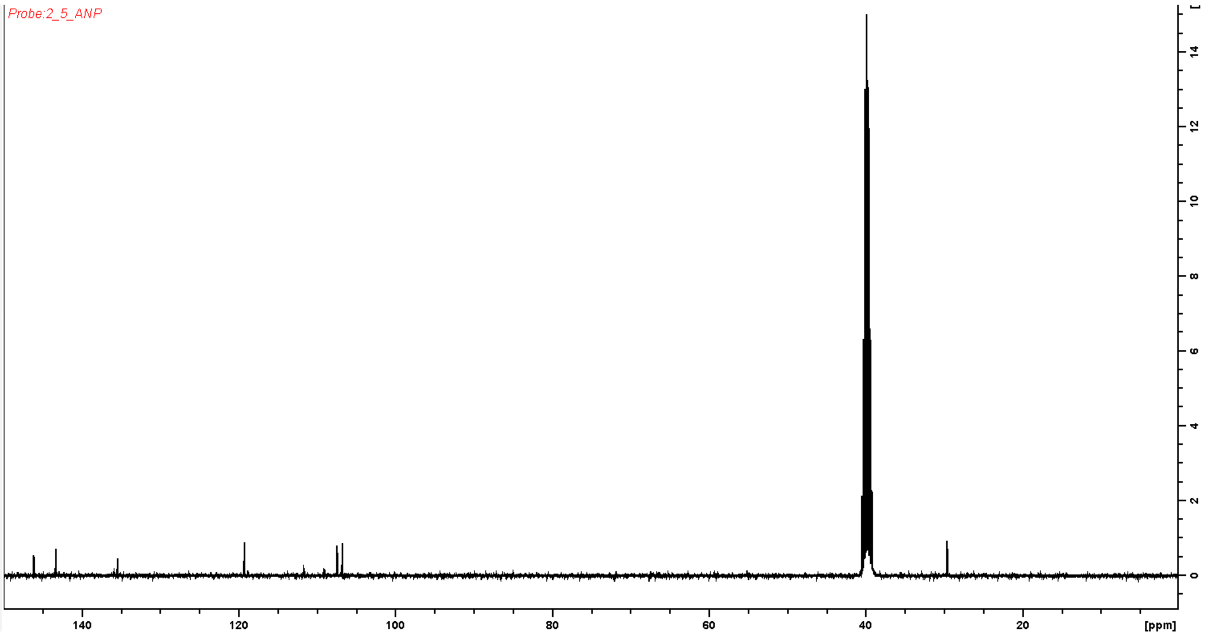

**Figure S 17** – $^{13}$C-NMR for product **4a** from in vivo upscale experiments with *Rg*ANMT after purification. $^{13}$C-NMR (100.6 MHz, $D_6$MSO) δ(ppm): 146.1, 143.2, 135.5, 119.4, 107.4, 106.8, 29.6. The signal at 29.6 was assigned to the carbon of the methyl group adjacent to the amino group.

#### in vitro

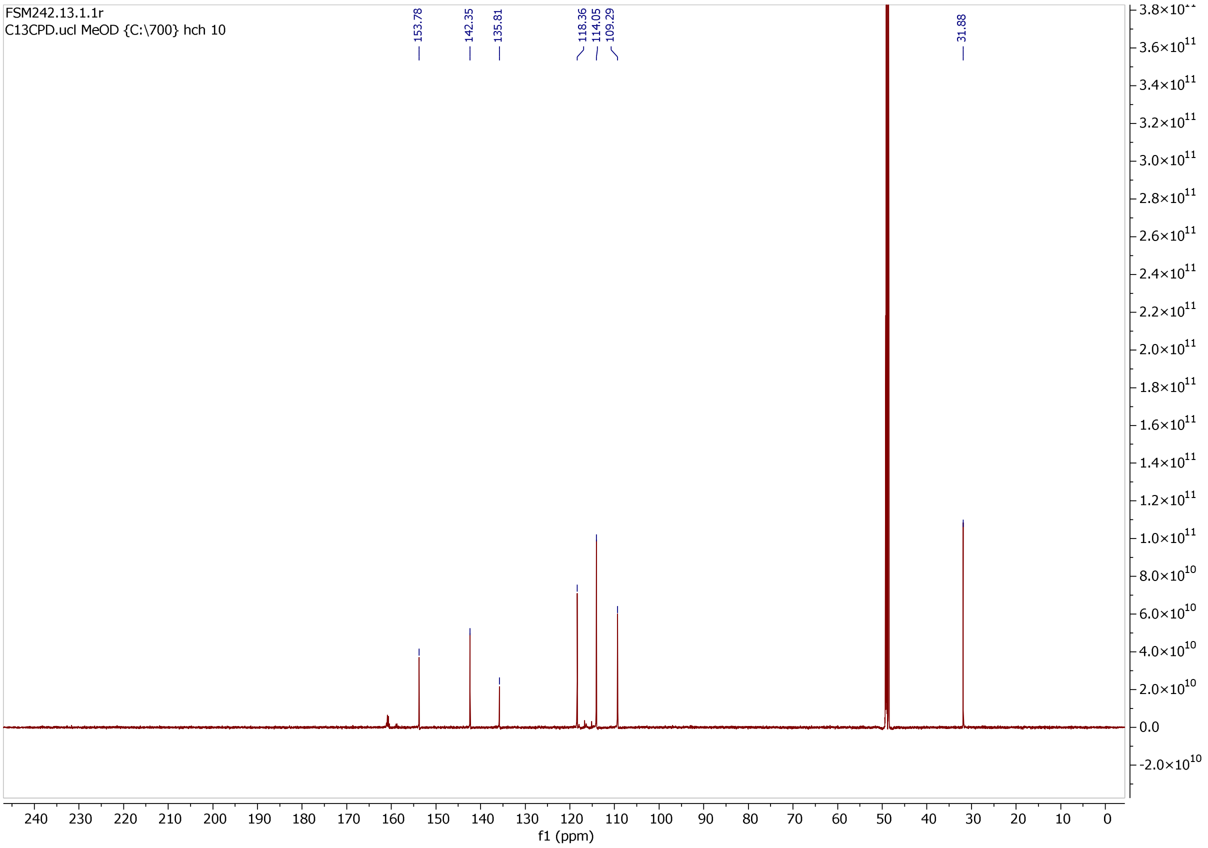

**Figure S 18** – $^{13}$C-NMR for product **3a** from in vitro upscale experiments with *Rg*ANMT after purification. $^{13}$C-NMR (176 MHz, MeOD) δ(ppm): 153.8, 142.4, 135.8, 118.4, 114.1, 109.9, 31.9. HRMS (ES+) found $[M+H]^+$ 169.0605; $C_7H_9N_2O_3$ requires 169.0608.

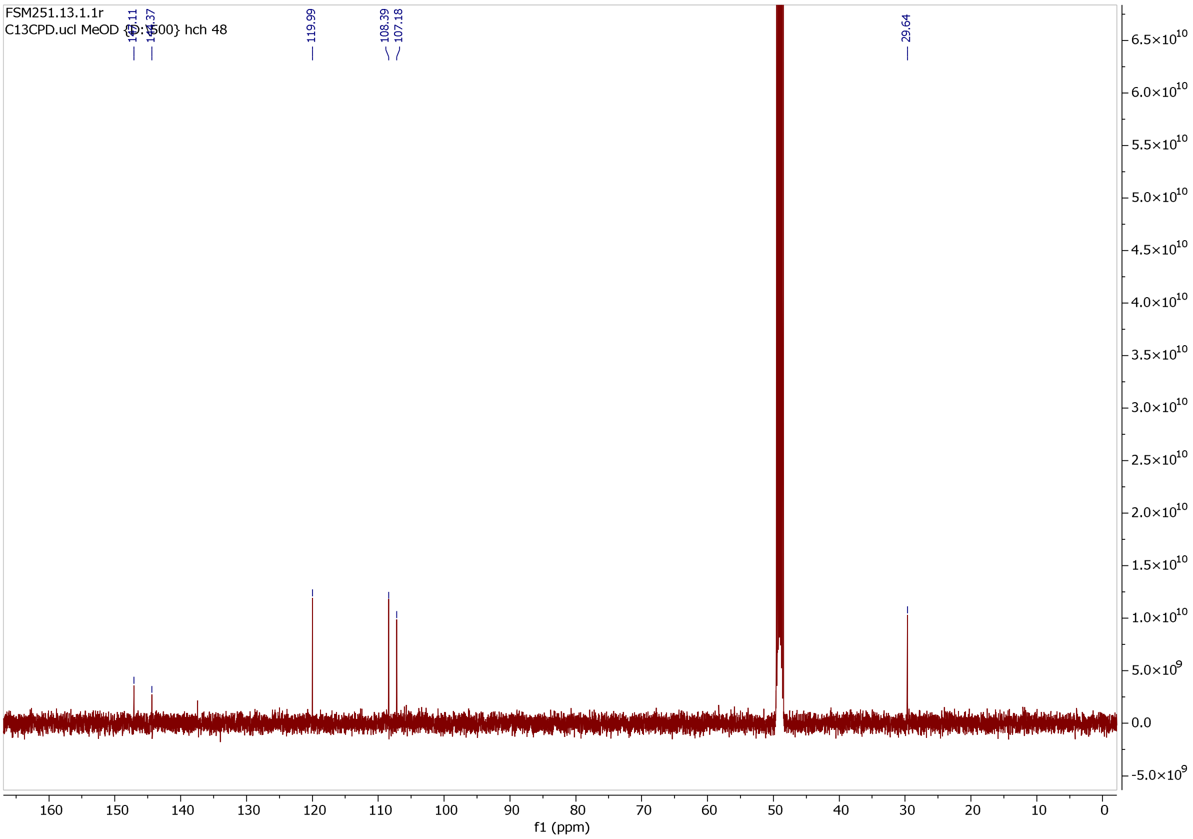

**Figure S 19** – $^{13}$C-NMR for product **4a** from in vitro upscale experiments catalysed by *Rg*ANMT after purification. $^{13}$C-NMR (126 MHz, MeOD) δ(ppm): 147.1, 144.4, 119.9, 108.4, 107.2, 29.6. HRMS (ES+) found $[M+H]^+$ 169.0607; $C_7H_9N_2O_3$ requires 169.06.

#### Synthesis *S*-allyl-L-homocysteine

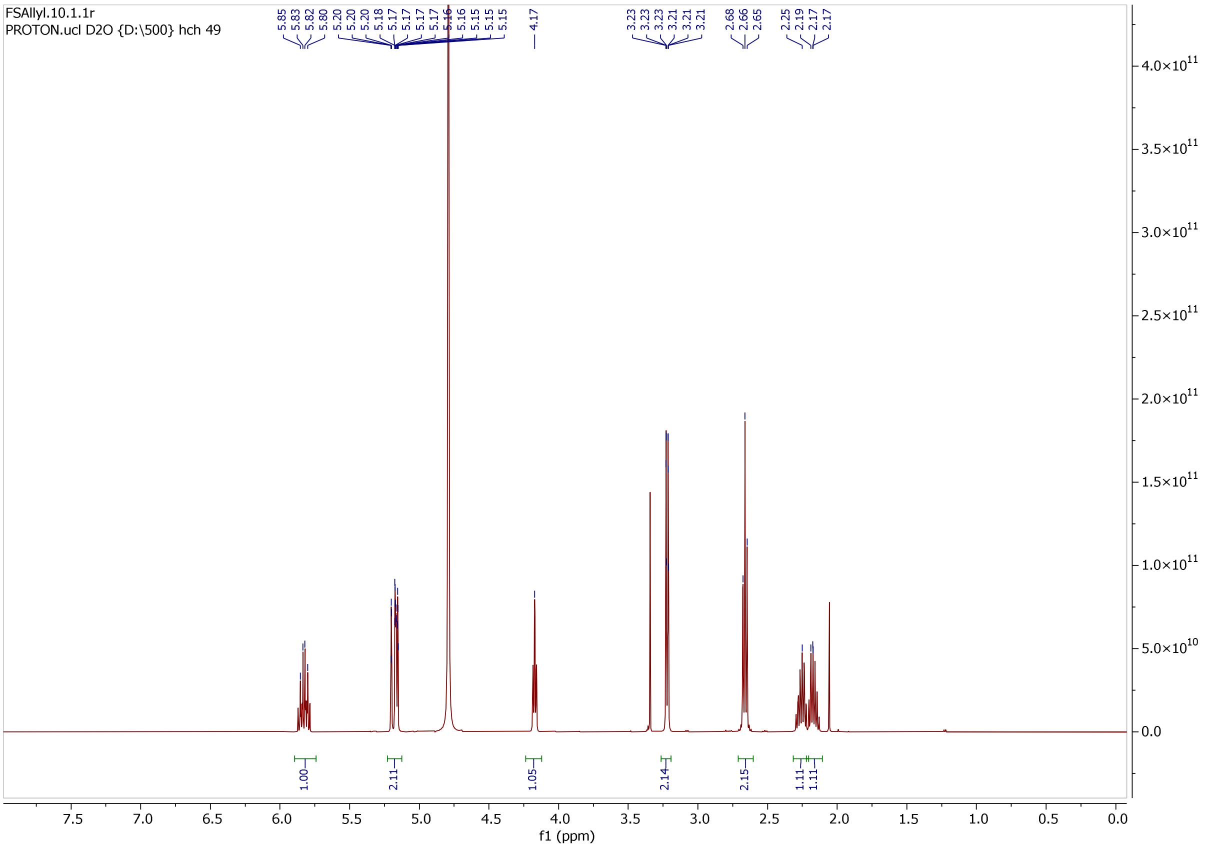

**Figure S 20** – $^{1}$H-NMR spectrum of chemical synthesis of S-allyl-L-homocysteine. $^{1}$H-NMR (500 MHz; $D_2O$) δ (ppm) = 2.25–2.17 (m, 2H), 2.68–2.65 (m, 2H), 3.23–3.21 (m, 2H), 4.17 (m, 1H), 5.20–5.15 (m, 2H), 5.85–5.80 (m, 1H).
